# The structure of the R2T complex reveals a different architecture of the related HSP90 co-chaperones R2T and R2TP

**DOI:** 10.1101/2024.03.27.587014

**Authors:** Alberto Palacios-Abella, Andrés López-Perrote, Jasminka Boskovic, Sandra Fonseca, Cristina Úrbez, Vicente Rubio, Oscar Llorca, David Alabadí

## Abstract

Heat shock protein 90 (HSP90) is a molecular chaperone that contributes to the maturation and activation of substrates in multiple cellular pathways. Its activity is supported by various co-chaperones. One of these is R2TP, a complex of RuvBL1-RuvBL2-RPAP3-PIH1D1 in humans, which is involved in the assembly of various multiprotein complexes, including mTORC1 and Box C/D and Box H/ACA snoRNPs. Structural analyses have shown that the complex is organized around a heterohexameric ring of the ATPases RuvBL1-RuvBL2 in both yeast and humans. In addition, several R2TP-like co-chaperones have been identified in humans, such as R2T, which lacks PIH1D1, but these are less well characterized. In seed plants, there are no PIH1D1 orthologs. Here, we have identified the R2T complex of *Arabidopsis* and determined its cryoEM structure. R2T associates with the prefoldin-like complex *in vivo* and is located in the cytosolic and nuclear compartments. R2T is organized as a dodecamer of AtRuvBL1-AtRuvBL2a that forms two rings, with one AtRPAP3 anchored to each ring. AtRPAP3 has no effect on the ATPase activity of AtRuvBL1-AtRuvBL2a and binds with a different stoichiometry than that described for human R2TP. We show the interaction of AtRPAP3 with AtRuvBL2a and AtHSP90 *in vivo* and describe the residues involved. Taken together, our results show that AtRPAP3 recruits AtRuvBL1-AtRuvBL2a and AtHSP90 via a mechanism that is also conserved in other eukaryotes, but that R2T and R2TP co-chaperone complexes have distinct structures that also suggest differences in their functions and mechanisms.

## INTRODUCTION

The chaperone HSP90 plays an important role in protein homeostasis in eukaryotes, especially under unfavorable cellular conditions when the activity of proteins and protein complexes is challenged (Schopf et al., 2017). Not surprisingly, the influence of HSP90 activity is ubiquitous in plants. For instance, HSP90 is necessary to buffer genetic variants that may have potential effects on protein stability or activity in *Arabidopsis* (Queitsch et al., 2002). Without adequate HSP90 activity, the phenotypic effects of such variants would otherwise manifest. In line with this, *Arabidopsis* plants with constitutively reduced HSP90 activity present altered development and circadian clock dysfunction (Kim et al., 2011), while a mutant lacking two of the four cytosolic HSP90 paralogs is lethal (Hubert et al., 2009).

HSP90 does not act alone but is supported by numerous co-chaperones that either drive the conformational changes associated with chaperone activity, act as adaptors to recruit client proteins, or both (Schopf et al., 2017). One of these co-chaperones is the R2TP complex, which supports HSP90 in the assembly of various protein complexes in yeast and mammals, e.g. the Box C/D and Box H/ACA small nucleolar ribonucleoproteins, which are involved in pre-rRNA modifications, or the TOR complex (Houry et al., 2018). R2TP stands for Rvb1-Rvb2-Tah1-Pih1, after these four proteins were identified as members of an HSP90-interacting complex in yeast (Zhao et al., 2005). Rvb1 and Rvb2 are AAA+ ATPases with orthologs in animals and plants named RuvB-like 1 (RuvBL1) and RuvBL2 (Figure 1A) (Kanemaki et al., 1997; Holt Iii et al., 2002; Nano and Houry, 2013; Majerska et al., 2017). Rvb1/RuvBL1 and Rvb2/RuvBL2 share > 40 % identity and have a similar structure (Dauden et al., 2021). Both proteins have an ATPase domain which is inserted by the prominent Domain II (DII) with a characteristic oligonucleotide/oligosaccharide binding-fold (OB-fold) responsible for protein-protein interactions. Rvb1/RuvBL1 and Rvb2/RuvBL2 are shared with other complexes involved in chromatin remodeling, such as SWR1 and INO80-C (Candela-Ferré et al., 2024). Yeast Tah1 is a small protein (12 KDa) consisting almost exclusively of a carboxylate clamp-type TPR domain that interacts with the C-terminal peptide MEEVD of HSP90 and is thus responsible for the recruitment of the chaperone to the R2TP complex (Zhao et al., 2005). In humans, Tah1 ortholog RPAP3 is a larger protein (75 KDa) containing two TPR domains and a specialized C-terminal domain named RuvBL2-Binding Domain (RBD) for being involved in the interaction with RuvBL2 (Jeronimo et al., 2007; Martino et al., 2018). The plant RPAP3 ortholog (44 KDa) containing a single TPR domain and the RBD has been identified in sorghum (Figure 1A) (Machado Antonio et al., 2023). Pih1 and its ortholog in animals PIH1D1 contain a C-terminal CS domain that mediates interaction with Tah1/RPAP3 and an N-terminal PIH domain that interacts with a specific phosphorylated motif in client proteins (Horejsi et al., 2014). Interestingly, a phylogenetic analysis has suggested that homologs of Pih1/PIH1D1 are exclusively found in bryophytes within the green lineage (Maurizy et al., 2018). Thus, the R2TP complex in vascular plants such as *Arabidopsis* would be formed by RuvBL1, RuvBL2 and RPAP3. In fact, this reduced version of the complex, called R2T, has been identified in humans in addition to the canonical one. The human R2T can be formed by RuvBL1, RuvBL2 and RPAP3-2, an RPAP3 isoform that lacks the PIH1D1-interacting motif (Maurizy et al., 2018), and by competition between RPAP3 and PIH1D1 for binding to RuvBL1-RuvBL2 (Wing et al., 2022). It is not known whether an R2T complex is formed *in vivo* in plants.

**Figure 1.**
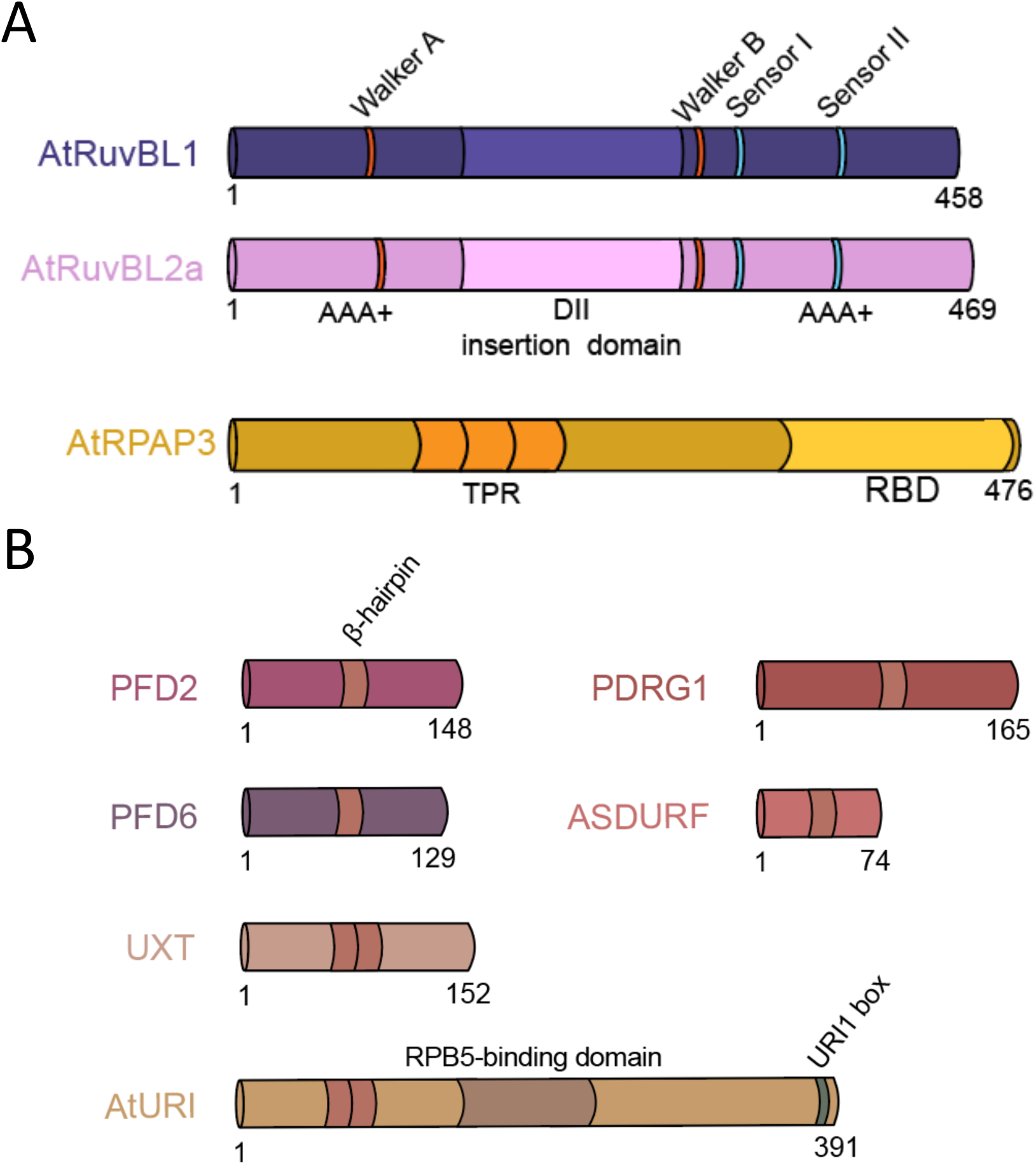
Schematic representation of the subunits of the *Arabidopsis* R2T and PFDL complexes. (A) Catalytic motifs involved in ATP hydrolysis, as well as the DII insertion domain and domains forming the AAA+ core, are indicated in AtRuvBL1 and AtRuvBL2a. The AtRPAP3 subunit contains a TPR motif and the RuvBL2 binding domain (RBD) at the C-terminus. (B) The PFD/PFDL schematics show the single and the double β-hairpin connecting the two helices in the α-type and β-type prefoldins, respectively. For AtURI, the RPB5-interacting domain and the conserved URI box are shown.

CryoEM analyses have provided structural insights into the 3D organization of *in vitro* assembled R2TP complexes from yeast and human (Rivera-Calzada et al., 2017; Martino et al., 2018; Muñoz-Hernández et al., 2019). The structure of the two complexes is similar in terms of the organization of the Rvb1/RuvBL1 and Rvb2/RuvBL2 proteins in a hetero-hexameric ring, built by alternating subunits, with the ATPase and DII domains occupying opposite sides of the ring. This defines two faces in each hetero-hexameric ring, the AAA-face containing the ATPase domains, and the DII-face at the opposite end of the complex. In yeast, Pih1 makes contacts with the Rvb1-RvB2 ring and with the C-terminal extension of Tah1, with the Pih1-Tah1 hetero-dimer housed in the basket formed by the DII domains of Rvb1-RvB2 (Rivera-Calzada et al., 2017). In yeast, the configuration of the R2TP suggests that the flexibility to bind clients is mainly provided by HSP90. In humans, RPAP3 plays a more important role in the organization of the complex than Tah1 in yeast. The C-terminal RBD of RPAP3 specifically anchors the protein to the ATPase side of the RuvBL1-RuvBL2 ring via an interaction with RuvBL2, while the rest of the protein, which is responsible for binding to PiH1D1 and HSP90, is flexible and thus may facilitate the recruitment of a variety of clients (Martino et al., 2018). Interestingly, each R2TP complex can accommodate up to three RPAP3 molecules per RuvBL1-RuvBL2 ring, one per each RuvBL2 (Martino et al., 2018). Although the human R2T version of the complex has been identified *in vivo*, it is not known how it is structurally organized in the absence of PIH1D1.

Several interactome studies have shown that the human R2TP is usually associated with the prefoldin-like complex (PFDLc) to form the R2TP/PFDL complex (Cloutier et al., 2009; Boulon et al., 2010; Cloutier et al., 2017; Cloutier et al., 2020), also known as PAQosome, for particle for arrangement of quaternary structure (Houry et al., 2018). The PFDLc is a hetero-hexameric complex consisting of the canonical prefoldins PFDN2 and PFDN6 and the PFDL proteins URI1, UXT, PDRG1 and ASDURF. Although the exact function of PFDLc is not known, it has been hypothesized that it may act as an adaptor to recruit clients into the complex, such as RNA polymerase II and other multi-subunit protein complexes (Houry et al., 2018). We have recently found that the PFDLc is present in plants (Gómez-Mínguez et al., 2024).

In this work, we have identified the *Arabidopsis* R2T complex. We found that R2T is associated with PFDLc *in vivo* and mediates HSP90 activity. Furthermore, we determined the 3D structure of the *in vitro* assembled R2T complex by cryoEM. Our results show that, in contrast to the R2TP, the R2T complex is preferentially organized as a dodecameric double ring of AtRuvBL1-AtRuvBL2a with an AtRPAP3 anchored to the ATPase side of either one or the two rings. The different architecture of the R2T and the related R2TP suggests that each of these co-chaperone complexes might have functional and mechanistic particularities.

## RESULTS

### The R2T complex is formed *in vivo* and associates to PFDLc in *Arabidopsis*

Studies in mammals have shown that the prefoldin PFDN6 is present in at least two alternative complexes: the canonical prefoldin complex (PFDc) (Gestaut et al., 2019) and the PFDLc (Gstaiger et al., 2003; Sardiu et al., 2008; Cloutier et al., 2020). Similarly, tandem affinity purification-mass spectrometry (TAP-MS) and AP-MS approaches have shown that the *Arabidopsis* PFDN6 ortholog, PFD6, is also part of both PFDc and PFDLc (Blanco-Touriñán et al., 2021; Gómez-Mínguez et al., 2024). To identify additional interactors of PFD6, we performed TAP-MS using *Arabidopsis* PSB-D cell suspensions (Van Leene et al., 2015). The GS^rhino^ tag, consisting of two protein G domains, followed by two repeats of the rhinovirus 3C protease cleavage site and the streptavidin-binding peptide, was fused to the N-terminus of PFD6 and the fusion protein was expressed in PSB-D cell suspensions. We identified numerous peptides corresponding to all subunits of the PFDL and canonical PFD complexes after performing the two consecutive purification steps (Figure 1B, Table 1 and Supplementary Figure S1), as we expected based on our previous results using TAP- or AP-MS with other subunits as baits (Blanco-Touriñán et al., 2021; Gómez-Mínguez et al., 2024). None of these proteins were included in the list of proteins that appear non-specific in TAP experiments with protein fusions to the GS^rhino^ tag (Van Leene et al., 2015). We also identified peptides from other proteins that are associated with PFDLc in humans (Cloutier et al., 2009; Cloutier et al., 2017; Cloutier et al., 2020). These include two paralog subunits present in all nuclear RNA polymerases, NRPB5A and NRPB5C (Supplementary Figure S2A), as well as components of the R2T complex (Table 1): AtRPAP3, AtRuvBL1, and AtRuvBL2a, one of the two AtRuvBL2 paralogs (Schorova et al., 2019). Although AtRuvBL1 and AtRuvBL2a appeared non-specifically in a series of TAP experiments (Van Leene et al., 2015), we considered them as true interactors given their close association with RPAP3 in the R2TP complex in other organisms (Martino et al., 2018; Maurizy et al., 2018; Wing et al., 2022).

**Table 1.** Subunits of the R2T and PFDL complexes identified by TAP-MS using GS-PFD6 as bait.

|  | AGI Code | Peptides<sup>a</sup> |  |
| --- | --- | --- | --- |
|  |  | R1 | R2 |
| AtRPAP3 | AT1G56440 | 2 | 5 |
| AtRuvBL1 | AT5G22330 | 7 | 8 |
| AtRuvBL2a | AT5G67630 | 4 | 10 |
| AtURI | AT1G03760 | 141 | 154 |
| UXT | AT1G26660 | 143 | 109 |
| PDRG1 | AT3G15351 | 48 | 44 |
| PFD2 | AT3G22480 | 437 | 491 |
| PFD6 | AT1G29990 | 669 | 697 |
| ASDURF | AT1G49245 | 18 | 23 |

<sup>a</sup>Number of total peptides of each protein identified in each replicate (R1-R2).

To confirm the presence of the R2T complex *in vivo* and its association with the PFDLc, we performed AP-MS in *Arabidopsis* PSB-D cell suspensions expressing GS^rhino^-AtRPAP3. We identified AtRuvBL1 and AtRuvBL2a as one of the major interactors (Table 2), which is consistent with the presence of the *Arabidopsis* R2T *in vivo*. We also identified all subunits of the PFDLc except ASDURF (Table 2). We found no canonical PFDs other than PFD2 and PFD6, suggesting that AtRPAP3 is unable to interact with the canonical PFDc as reported in humans (Cloutier et al., 2009). Overall, our results show that the R2T complex is formed in *Arabidopsis* and that it associates with the PFDLc.

**Table 2.** Subunits of the R2T and PFDL complexes identified by AP-MS using GS-AtRPAP3 as bait.

|  | AGI code | Peptides<sup>a</sup> |  |  | Control<sup>b</sup> |  |
| --- | --- | --- | --- | --- | --- | --- |
|  |  | R1 | R2 | R3 | R1 | R2 |
| AtRPAP3 | AT1G56440 | 106 | 89 | 69 | 1 | 3 |
| AtRuvBL1 | AT5G22330 | 82 | 67 | 44 | 15 | 19 |
| AtRuvBL2a | AT5G67630 | 94 | 70 | 53 | 17 | 23 |
| AtURI | AT1G03760 | 26 | 19 | 17 | 0 | 0 |
| UXT | AT1G26660 | 5 | 1 | 0 | 0 | 0 |
| PDRG1 | AT3G15351 | 4 | 4 | 2 | 0 | 0 |
| PFD2 | AT3G22480 | 8 | 9 | 6 | 0 | 0 |
| PFD6 | AT1G29990 | 0 | 2 | 4 | 0 | 0 |
| ASDURF | AT1G49245 | 0 | 0 | 0 | 0 | 0 |

<sup>a</sup>Number of total peptides of each protein identified in each replicate (R1-R3).
<sup>b</sup>Number of total peptides of each protein identified in the control purification (R1-R2).

### Direct protein-protein interactions within the R2T complex

We determined the direct interactions between the subunits of the R2T complex using yeast two-hybrid (Y2H) assays. As expected based on previous results in *Arabidopsis* (Schorova et al., 2019), we observed interactions between AtRuvBL1 and AtRuvBL2a (Figure 2 and Supplementary Figure S3), suggesting that these two proteins likely form hetero-multimers, as in other eukaryotes (Dauden et al., 2021). The similar behavior of these proteins is consistent with the high degree of sequence homology between *Arabidopsis* and human orthologs (Supplementary Figure S4). Next, we tested the direct interactions between these two subunits and AtRPAP3. AtRuvBL2a was able to interact with AtRPAP3, whereas we did not observe any interaction of the latter with AtRuvBL1 (Figure 2 and Supplementary Figure S3). This interaction pattern corresponds to that of the human R2TP complex, in which RPAP3 comes into direct contact with RuvBL2 (Martino et al., 2018; Maurizy et al., 2018). Although the degree of homology between *Arabidopsis* and human RPAP3 is relatively low compared to the orthologs of RuvBL1 and RuvBL2 in both species (Supplementary Figure S4), their ability to interact specifically with RuvBL2 is conserved.

**Figure 2.**
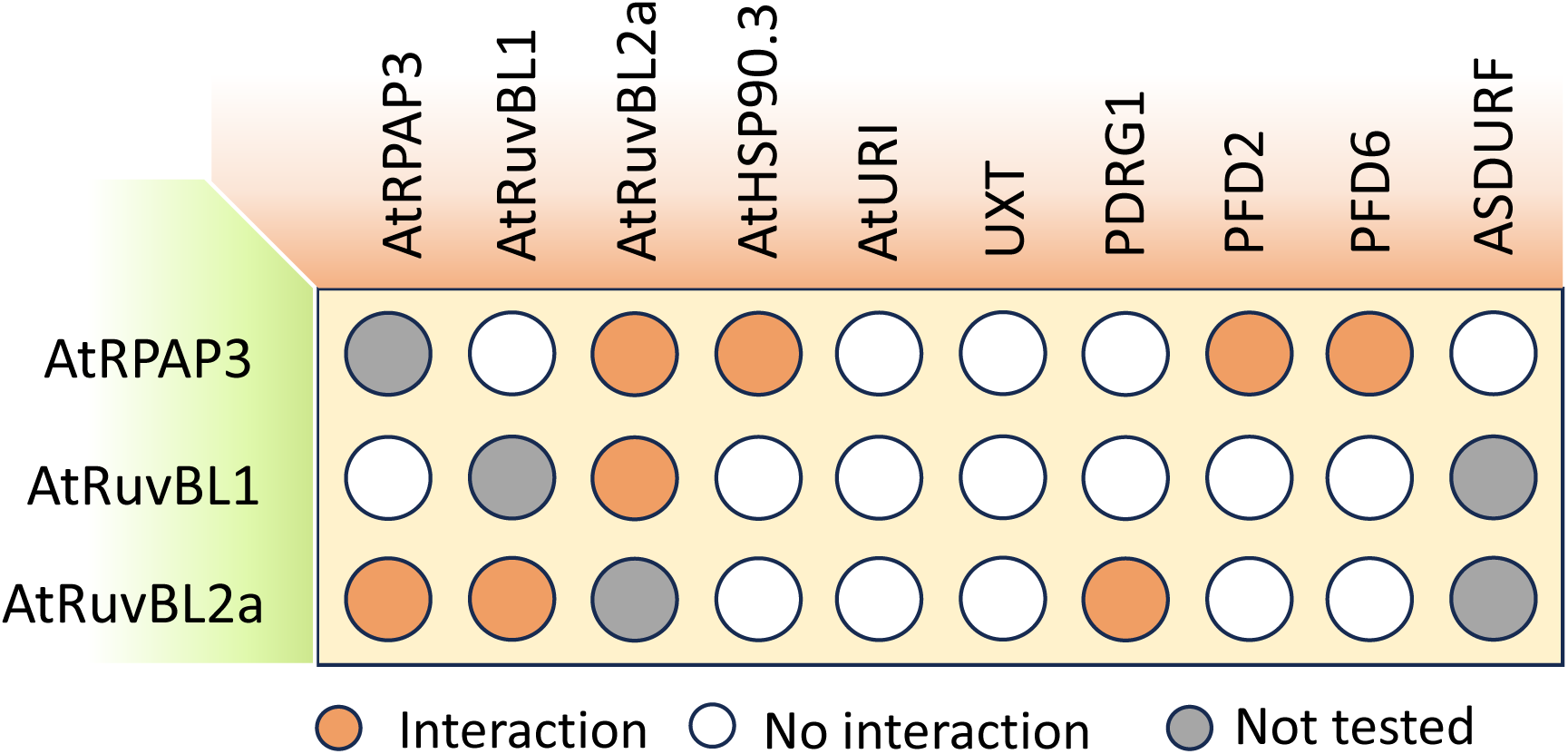
Direct interactions between subunits of R2T and PFDL complexes. Diagram showing the results of the Y2H assays of the interactions between the subunits of the R2T and PFDL complexes.

Next, we confirmed the interaction between AtRPAP3 and AtRuvBL2a and indirectly with AtRuvBL1 by pull-down experiments with recombinant proteins expressed in *E. coli*. We fused the His-tag and the solubility partner IF2DI to the N-terminal end of AtRPAP3 and the Strep-tag to its C-terminus. AtRPAP3 was co-expressed with His-AtRuvBL1 and untagged AtRuvBL2a. We were able to co-purify AtRuvBL1 and AtRuvBL2a together, in an AtRPAP3-dependent manner, when we pulled down AtRPAP3 via the Strep-tag (Figure 3A). These results indicate that the three proteins form the R2T complex. Similarly, *in vitro* pull-down assays showed that sorghum SbRPAP3 can form a complex with human RuvBL1 and RuvBL2 (Machado Antonio et al., 2023).

**Figure 3.**
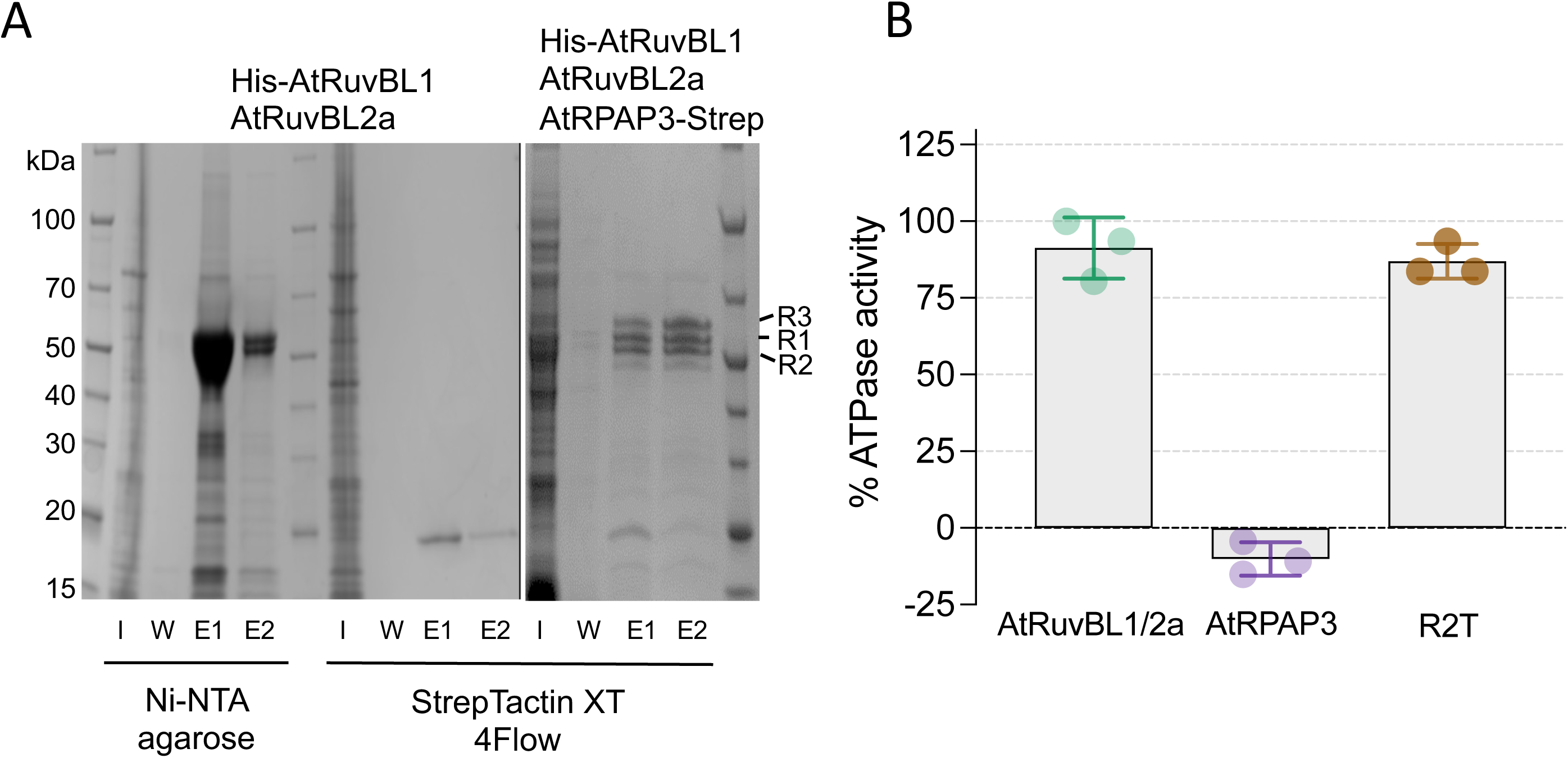
Assembly of R2T *in vitro* and ATPase activity. (A) Pull-down assays. Control experiment of AtRuvBL1-AtRuvBL2 purification using Ni-NTA and StrepTactin XT resins (left). An *E. coli* culture expressing AtRuvBL1-AtRuvBL2a was split in two after lysis and clarification, and incubated with Ni-NTA agarose resin or StrepTactin XT resin. After extensive washing, proteins were eluted with the appropriate buffer (imidazole in the case of Ni-NTA resin, and biotin for the StrepTactin XT resin) and analyzed by SDS-PAGE and Coomassie staining. Expression of AtRuvBL1-AtRuvBL2a was corroborated in the elution from the Ni-NTA resin, were both proteins co-eluted (left). An *E. coli* culture expressing AtRPAP3 (R3)-AtRuvBL1 (R1)-AtRuvBL2a (R2) was lysed, clarified and incubated with StrepTactin XT resin (right). After extensive washing, proteins were eluted with biotin and analyzed by SDS-PAGE and Coomassie staining. AtRuvBL1-AtRuvL2a were eluted from the StrepTactin XT resin, indicating binding to AtRPAP3 (right). Input (I), wash (W), elution 1 (E1) and elution 2 (E2) for both experiments are shown. (B) ATPase activity assays of AtRuvBL1-AtRuvBL2a in absence and presence of AtRPAP3. Control experiment of AtRPAP3 alone is shown. Relative ATPase activities are represented as mean ± S.D. values from three replicates, considering ATP hydrolysis by AtRuvBL1-AtRuvBL2a is considered as 100% activity.

### AtRPAP3 does not contribute to the ATPase activity of the R2T complex

As shown above, AtRPAP3 interacts with AtRuvBL1, AtRuvBL2a to assemble the R2T complex (Figure 3A). Then, we investigated if the interaction of AtRPAP3 with AtRuvBL1-AtRuvBL2a would have any effect on the ATPase activity of AtRuvBL1-AtRuvBL2a. For this, we purified to homogeneity AtRuvBL1-AtRuvBL2a and AtRPAP3 (Supplementary Figures S5A and S5B) and then we performed time course measurements to monitor de ATP consumption by the ATPases in the absence and presence of AtRPAP3 (Figure 3B). An average rate of 9.33 ± 1.01 min^-1^ of ATP turnover was calculated for AtRuvBL1-AtRuvBL2a, similar to what have been described for the human orthologs (López-Perrote et al., 2020), whereas AtRPAP3 alone did not exert measurable ATP hydrolysis. Addition of AtRPAP3 to AtRuvBL1-AtRuvBL2a did not significantly affect the catalytic activity of the proteins (8.88 ± 0.57 min^-1^ of ATP turnover), suggesting that AtRPAP3 is not involved in the regulation of the AtRuvBL1-AtRuvBL2a ATPase activity. These results are consistent with those obtained in humans, where the ATPase activity of R2T was very similar to that of the RuvBL1-RuvBL2 hetero-hexamer (Wing et al., 2022).

### Mapping direct protein-protein interactions of R2T with HSP90 and PFDLc

One of the functions of RPAP3 within the R2TP complex in animals and yeast is the recruitment of the chaperone HSP90 via the interaction between the conserved C-terminal end of the chaperone and the TPR domain of RPAP3 (Pal et al., 2014). The TPR domain of AtRPAP3 shows significant homology with human RPAP3 TPR2 that specifically interacts with HSP90 (Supplemental Figure 4), as observed for sorghum SbRPAP3 (Machado Antonio et al., 2023). We found that AtRPAP3 was able to interact with AtHSP90.3, one of the four highly homologous cytosolic HSP90 isoforms in *Arabidopsis* (Krishna and Gloor, 2001), by Y2H (Figure 2 and Supplemental Figure 3), suggesting that the role of RPAP3 in the recruitment of HSP90 is conserved in plants.

Since we have identified five of the six subunits of the PFDLc as interaction partners of RPAP3, we next determined which subunits make contacts between the R2T and PFDL complexes. The Y2H analysis showed that AtRuvBL2a can interact with PDRG1, whereas AtRPAP3 does so with PFD2 and PFD6 (Figure 2 and Supplementary Figure S3). This result shows that several subunits are involved in establishing contacts between the two complexes, indicating the potential for a close association.

### The R2T complex localizes in both the cytoplasm and the nucleus in *Arabidopsis*

The analysis of normalized protein accumulation in various *Arabidopsis* organs using ProteomicsDB (Samaras et al., 2020) revealed overlapping patterns for AtRPAP3, AtRuvBL1, and AtRuvBL2a, consistent with their participation in the same complex (Supplementary Figure S6A). Next, we determined the subcellular location where the R2T complex is formed. YFP-AtRPAP3 was localized to both the nucleus and cytoplasm when transiently expressed in *Nicotiana benthamiana* leaves as either YFP or RFP fusion (Figure 4A and Supplementary Figure S6B), consistent with its localization in *Arabidopsis* (Sotta et al., 2016). A very similar localization was observed for YFP-AtRuvBL2a in leaves of *N. bethamiana*, as expected for proteins acting in the same complex (Figure 4B). To confirm this, we performed Bimolecular Fluorescence Complementation (BiFC) by transient expression of YFN-AtRPAP3 and YFC-AtRuvBL2a in leaves of *N. benthamiana*. We observed the fluorescence of the reconstituted YFP in both the cytoplasm and nucleus, while the YFP fluorescence of the negative control pairs was below the detection limit (Figure 4C). These results indicate that AtRPAP3 and AtRuvBL2a can interact at both subcellular locations, likely reflecting the localization of the R2T complex. This result is in contrast to the R2T complex in animals, which is assumed to perform nuclear functions due to the preferential localization of the RPAP3-2 isoform in the nucleus (Maurizy et al., 2018).

**Figure 4.**
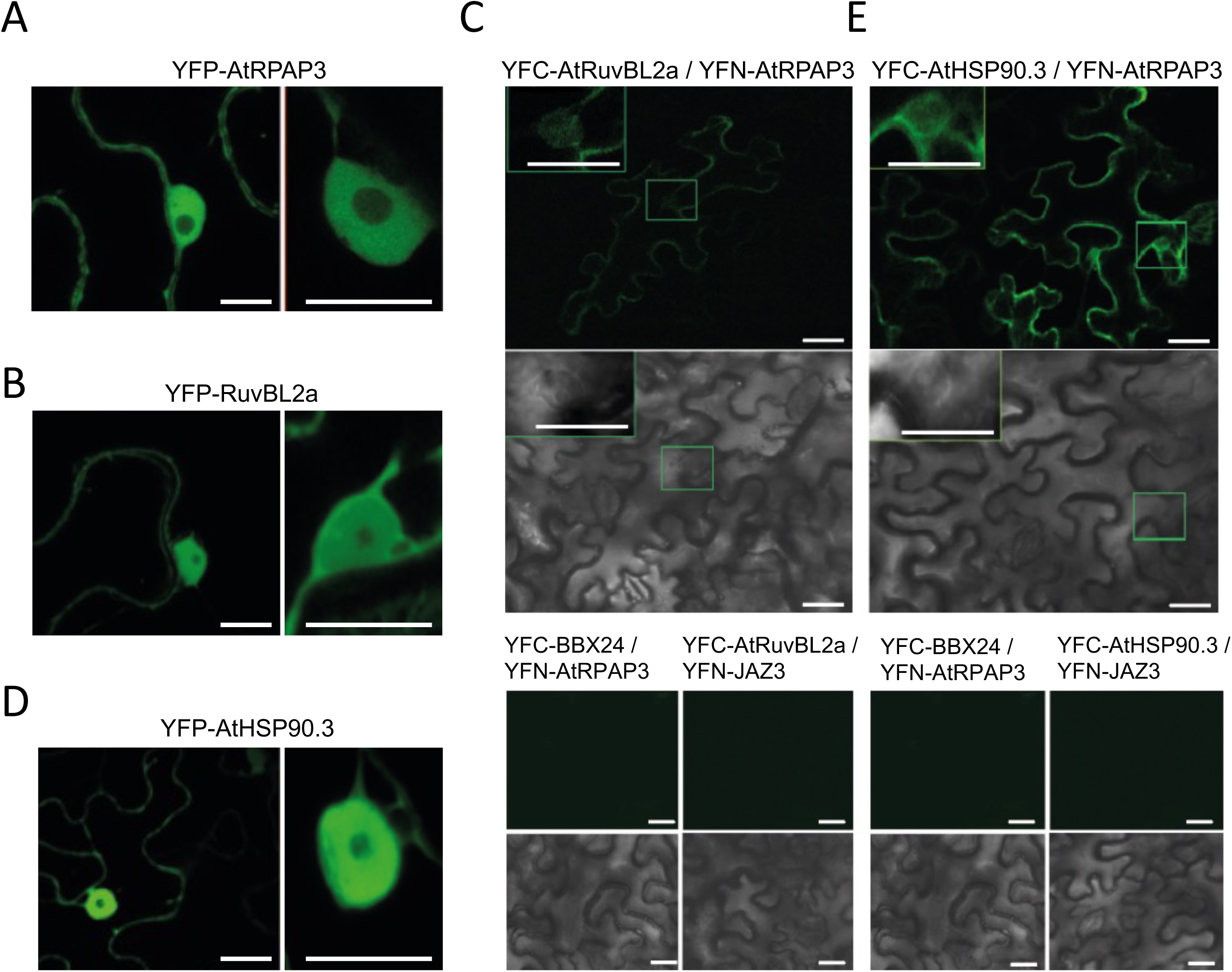
Subcellular localization of AtRPAP3, AtRuvBL2a and AtHSP90.3 in *N. benthamiana* and their interaction. (A, B, D) Nuclear and cytoplasmic localization of AtRPAP3, AtRuvBL2a and AtHSP90.3, respectively (scale bar: 10 μm). (C, E) BiFC assays in leaves of *N. benthamiana*. AtRPAP3 fused to the N-terminal YFP tag (YFN) was co-expressed with AtRuvBL2a (C) and AtHSP90.3 (E) fused to the C-terminal YFP tag (YFC). Fluorescence of the reconstituted YFP was detected by confocal microscopy of leaf disks three days after infiltration. YFN-AtJAZ3 and YFC-AtBBX24, which do not interact with AtRPAP3, AtRuvBL2a and AtHSP90.3, were used as negative controls. The insets show the fluorescence of a representative nucleus. Bright field images are shown below. Scale bar: 25 μm.

Consistent with the fact that the R2T complex also functions as a cochaperone of HSP90 in plants, YFP-AtHSP90.3 showed similar subcellular localization to AtRPAP3 and AtRuvBL2a when expressed in leaves of *N. benthamiana* (Figure 4D). Similarly, BiFC analysis showed that the interaction between AtRPAP3 and AtHSP90.3 occurs in the same subcellular localizations as that of AtRPAP3 and AtRuvBL2a (Figure 4E). Overall, these results are consistent with the idea that the R2T complex is formed in plants and that it can function as a cochaperone of HSP90.

### CryoEM of the *Arabidopsis* AtRuvBL1-AtRuvBL2a complex

The lack of an ortholog of PIH1D1 in *Arabidopsis* suggests that the overall structure of the R2T complex could be different from that of the R2TP. Proteomic and biochemical approaches have shown that an R2T complex can be formed in humans (Maurizy et al., 2018; Wing et al., 2022). However, the structure of the R2T complex is currently unknown. Therefore, we attempted to determine the structure of the R2T complex of *Arabidopsis*.

We first analyzed the structure of the AtRuvBL1-AtRuvBL2a complex using cryoEM. AtRuvBL1-AtRuvBL2a was purified to homogeneity after recombinant expression as a complex containing equimolar amounts of each protein and then analyzed by cryoEM (Supplementary Figures S5A, S5C and S5D). Images of single molecules of the complex in the cryoEM movies were extracted and subjected to image processing (Supplementary Figure S5C). Most of the reference-free 2D averages resembled those of human RuvBL1-RuvBL2, corresponding to the ring-shaped top views of AtRuvBL1-AtRuvBL2a and the barrel-like side views of dodecameric complexes containing 2 hetero-hexameric rings (Supplementary Figure S5D). We also found a minor population of side views corresponding to single ring hexameric complexes. Low resolution 3D reconstruction of the dodecameric complex showed two hexameric rings interacting back-to-back through the DII domains (Supplementary Figure S5E), similar to what has been observed for the highly homologous human RuvBL1-RuvBL2 complex (Gorynia et al., 2011). The two rings in the complex are flexibly attached, and thus we applied a focus refinement protocol on one of the rings that improved the resolution of the cryoEM map for this section of the complex to 4.7 Å (Supplementary Figure S5E). This allowed the fitting of the atomic structure of the human AAA core of the proteins (PDB 2XSZ) (Gorynia et al., 2011) into the cryoEM map, showing a high degree of similarity between the two species at this level of resolution (Supplementary Figure S5E, right-hand panel).

### CryoEM of the *Arabidopsis* R2T complex

We purified the R2T complex of *Arabidopsis* in amounts and purity suitable for structural studies (Supplementary Figure S7A). CryoEM and image processing of the R2T of *Arabidopsis* revealed molecules comprising the dodecameric form of the AtRuvBL1-AtRuvBL2a complex (Supplementary Figure S7B, Supplementary Table S1). This contrasts with the human R2TP, which consists of a hetero-hexameric RuvBL1-RuvBL2 ring, as PIH1D1, which is not present in *Arabidopsis*, disrupts the interaction between the two RuvBL1-RuvBL2 rings found in the dodecamers. In addition, some faint densities were observed at the AAA side of each AtRuvBL1-AtRuvBL2a ring, which we attributed to the binding of AtRPAP3 (Supplementary Figure S7B, yellow arrows in the right panel). Based on the structural information of the human R2TP complex, these densities could correspond to the C-terminal RBD of AtRPAP3 (Martino et al., 2018). The density of the TPR domain and the flexible region of AtRPAP3 was not visible in the cryoEM images. We hypothesize that this is due to the high flexibility of this region, similar to that described for the human complex (Martino et al., 2018).

In human R2TP, the RuvBL1-RuvBL2 ring can be decorated by up to 3 RBDs, one binding to each RuvBL2 subunit. Thus, the cryoEM was then processed using a workflow specifically designed to characterize the occupancy of the RBDs in each AtRuvBL1-AtRuvBL2a ring (Supplementary Figure S8). First, we implemented a 3D classification strategy focused on defining the occupancy of one of the rings. We found that 61.6 % of the particles contained no RBD (free AtRuvBL1-AtRuvBL2a), while 38.4 % contained one (R2T) (Supplementary Figure S8). Free AtRuvBL1-AtRuvBL2a may reflect disassembly of the complex during preparation of cryoEM samples, as purification was performed by pulling down from the AtRPAP3 subunit. Particles containing the RBD were realigned based on the position of this domain and then the occupancy of the opposite ring was analyzed in a similar manner. After selecting the best quality particles from a 3D classification based on this ring, further processing revealed the co-existence of two populations of R2T complexes. Complex 1 contained one RBD per ring (18% of the particles), while in complex 2 (82% of the particles) only one of the rings was bound to an RBD (Figure 5A and Supplementary Figure S8). Unexpectedly, the RBD in each ring of R2T complex 1 occupied a defined relative position, and we were unable to find any R2T complex that for a given position of an RDB in one of the rings would show an alternative position of the RBD in the opposite ring. This is a strong indication of some form of communication between the two rings in the dodecamer.

**Figure 5.**
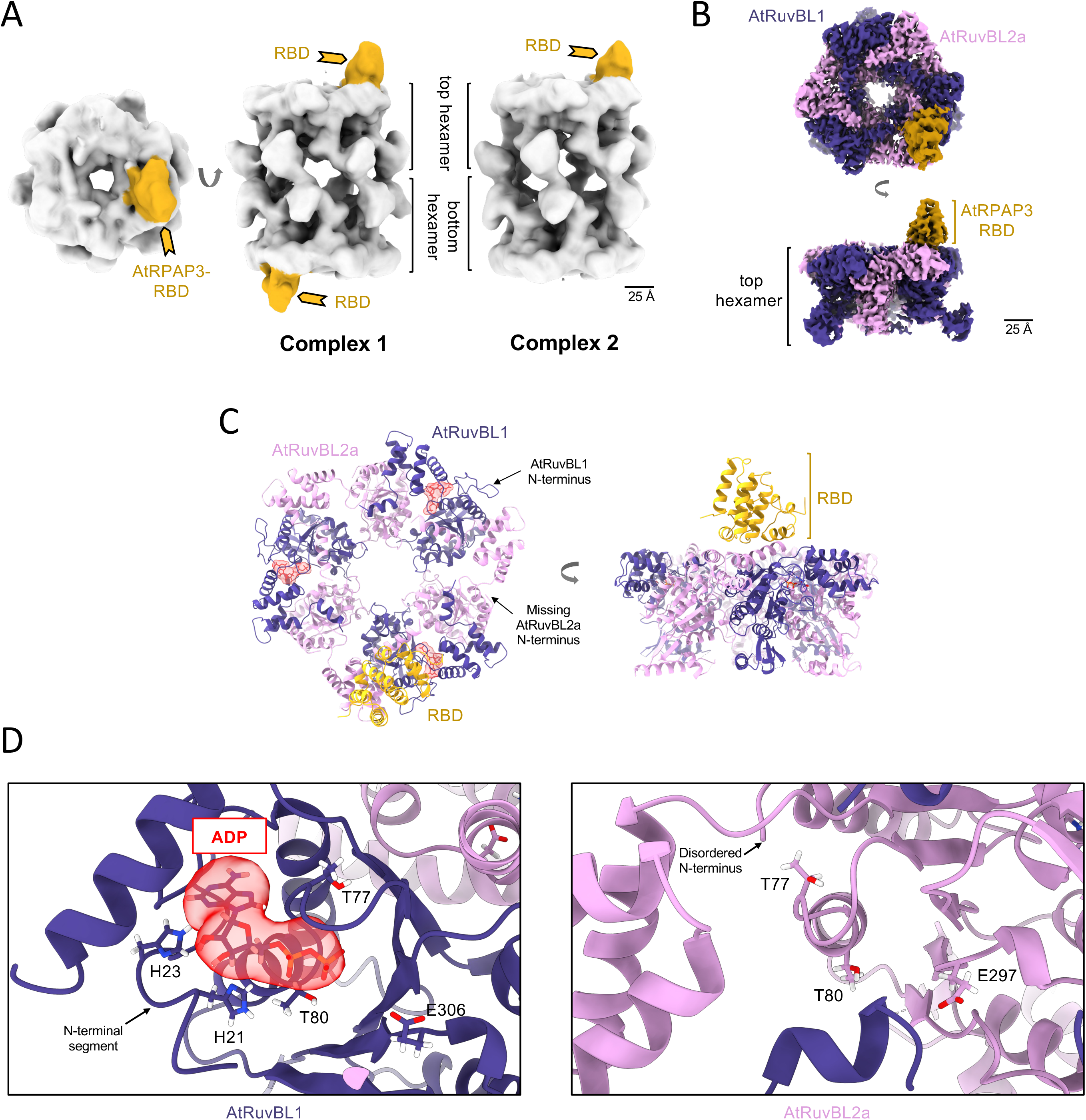
Architecture of the *Arabidopsis* R2T complex. (A) Low resolution structures of the dodecameric R2T complexes. The complex displayed a high degree of conformation flexibility between the two stacked hexameric rings. Implementation of 3D classification methods allowed the identification of two (complex 1) or one (complex 2) molecules of AtRPAP3 bound to the AAA face of the hexamers. Note that only the RBD domain of AtRPAP3 (in yellow) was visualized due to the flexibility of the TPR and linker regions of the protein. Interaction of the RBD domains was orchestrated in an organized manner in opposite sides of the AtRuvBL1-AtRuvBL2a dodecamer. (B) High resolution cryoEM map of the top ring of R2T complex obtained by focus refinement of the full dodecameric complex. (C) Atomic model for the top hexamer R2T complex. Nucleotide binding pocket of AtRuvBL1 subunits were filled with ADP (red transparent density), while AtRuvBL2a subunits were in an apo-state with no nucleotide bound. (D) Close-up views of the nucleotide binding pockets of AtRuvBL1 (left) and AtRuvBL2a (right). Atomic models of representative subunits are shown highlighting some catalytic residues. CryoEM density for the ADP molecule observed in AtRuvBL1 subunits is shown in red transparency with the atomic model of the nucleotide inside. The nucleotide binding pockets of the AtRuvBL2a subunits where empty and the N-terminal region disordered.

### Structure of the AtRuvBL1-AtRuvBL2a-RBD interaction

To gain insight into the interaction between AtRuvBL1-AtRuvBL2a and AtRPAP3 in the R2T complex, we conducted a focus 3D refinement on one of the AtRuvBL1-AtRuvBL2a rings containing an RBD to improve resolution. The resolution of the cryoEM map obtained (3.75 Å) allowed atomic model building of the structure for the interaction between AtRuvBL1-AtRuvBL2a and the RBD (Figures 5B and 5C and Supplementary Figures S9A and S9B). The DII domains were excluded from this analysis because their flexibility did not allow us to reach sufficient resolution for modeling.

The AAA ring of the R2T complex displays a similar structure to the human R2TP complex (Martino et al., 2018), with the RBD of AtRPAP3 attached to the AAA-face through the interaction with the AtRuvBL2a subunit of the hetero-hexameric AtRuvBL1-AtRuvBL2a ring (Figures 5A-C). However, this time we only found 1 RBD per ring compared to human R2TP, although the reconstitution strategy used for each complex was different (Martino et al., 2018). The AtRuvBL1 subunits were occupied by ADP and their N-terminal end, containing two conserved histidine residues, contributed to the binding of ADP trapped in the nucleotide binding pocket (Figure 5D). However, all AtRuvBL2a subunits were in an apo state with no bound nucleotide, which correlated with the lack of density for the N-terminal region of each AtRuvBL2a subunit in the cryoEM map, suggesting that they are disordered (Figure 5D). The N-terminal region of RuvBL2 has been described as a gate-keeper segment that controls nucleotide status in several human RuvBL1-RuvBL2-containing complexes and may represent a mechanism for regulating nucleotide exchange and/or hydrolysis (Muñoz-Hernández et al., 2019; López-Perrote et al., 2020; Serna et al., 2021).

The atomic structure of AtRPAP3 RBD comprises 8 helical segments (α1-α8), similar to the human orthologue (Supplementary Figures S9C and S9D). The RBD interacts with the helices α1, which is in contact with the C-terminal region of AtRuvBL1, as well as with α6 and the loop connecting α5-α6, which interact with a positively charged region of AtRuvBL2a (Figure 6A). Inspection of the interaction surface revealed that AtRPAP3 residues R428 and M431 make contacts with AtRuvBL2a. It should be noted that these two residues are conserved in humans (Supplementary Figure S4; marked with green asterisks), where their mutation impairs the interaction between RPAP3 and RuvBL2 (Maurizy et al., 2018).

**Figure 6.**
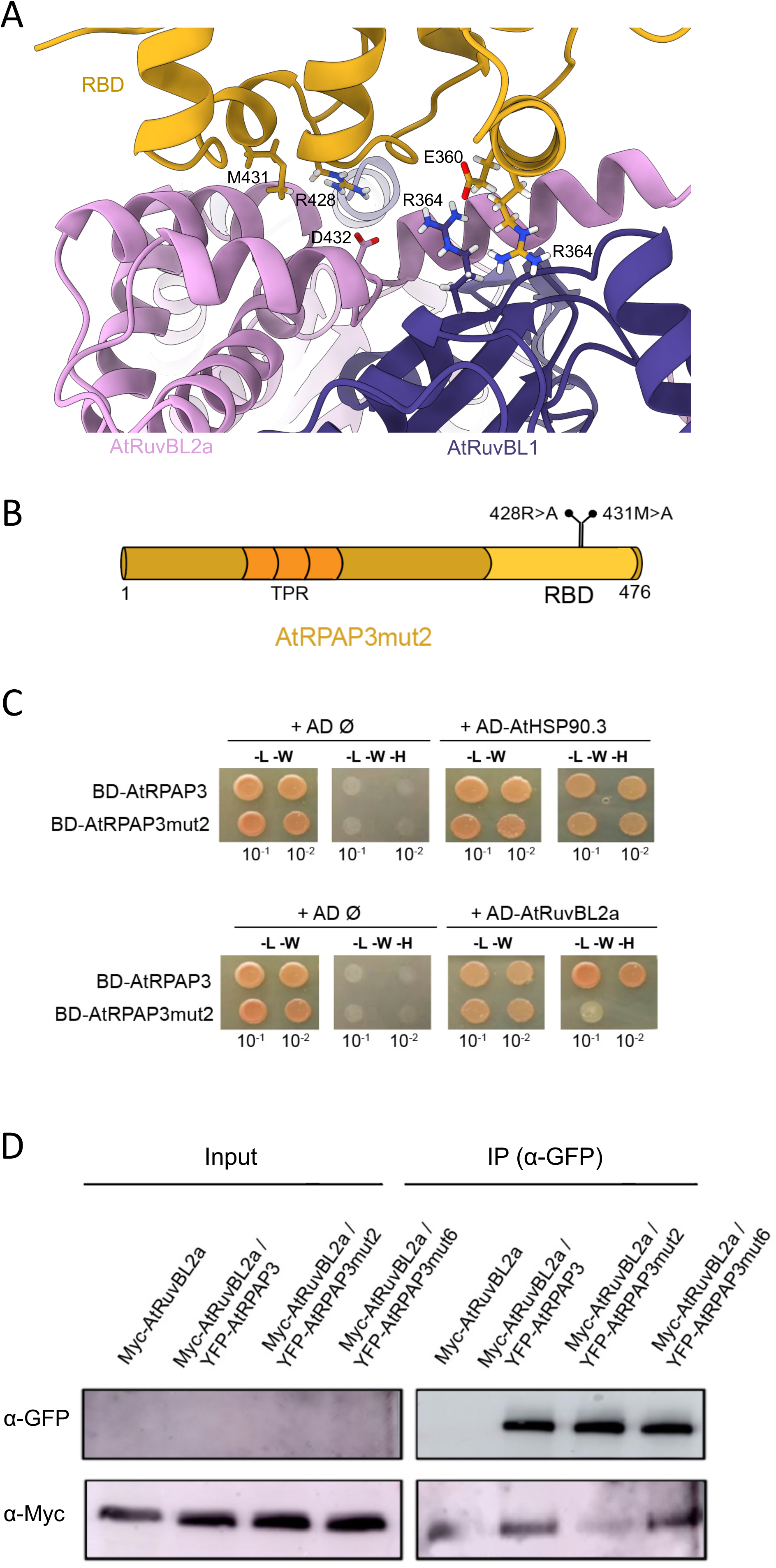
Mutations R428A and M431A in AtRPAP3 impair the AtRPAP3-AtRuvBL2a interaction *in vivo*. (A) Atomic model identifying the residues involved in the interaction between AtRuvBL2a and the RBD of AtRPAP3. (B) Schematic of AtRPAP3mut2, a mutant version of AtRPAP3, indicating the amino acid changes in the RBD domain. (C) Y2H assays for the interaction between AtRPAP3 and AtRPAP3mut2, both fused to the GAL4-BD, and AtRuVBL2a and AtHSP90.3, both fused to the GAL4-AD. L, leucine; W, tryptophan; H, histidine. The numbers indicate the dilution used for the assay. (D) Co-immunoprecipitation assay showing the interaction between AtRPAP3, AtRPAP3mut2 and AtRPAP3mut6 fused to YFP, and AtRuvBL2a fused to a Myc tag in *N. benthamiana*. Total proteins from the leaf extracts were immunoprecipitated with anti-GFP antibody-coated magnetic beads. The proteins were detected with anti-GFP and anti-Myc antibodies.

### The mutations R428A and M431A in AtRPAP3 impair the AtRPAP3-AtRuvBL2a interaction *in vivo*

To determine the importance of R428 and M431 for interaction *in vivo*, we used a mutant version of AtRPAP3 in which these two residues were changed to A (Figure 6B), as this change in human RPAP3 prevents interaction with RuvBL2 (Maurizy et al., 2018). A Y2H analysis showed that indeed the mutant version of AtRPAP3 (AtRPAP3mut2) has lost the ability to interact with AtRuvBL2a (Figure 6C, bottom panel). The lack of interaction was not due to an overall negative impact of the double mutation on AtRPAP3mut2, as it retained the ability to interact with AtHSP90.3 (Figure 6C; see also Figure 7D). To confirm that the mutations also prevent interaction *in planta*, we tested the interaction of myc-AtRuvBL2a with the two versions of YFP-AtRPAP3 by co-immunoprecipitation (co-IP) after transient expression in leaves of *N. benthamiana*. The different versions of YFP-AtRPAP3 were difficult to extract with the co-IP buffer and the protein signal after immunoblotting was below the limit of detection in the lanes corresponding to the inputs (Figure 6D). However, the proteins were visible after IP with an anti-GFP antibody. Importantly, myc-AtRuvBL2a was efficiently immunoprecipitated with anti-GFP antibodies from extracts co-expressing YFP-AtRPAP3 or the unrelated mutant YFP-AtRPAP3mut6 (see below), whereas the interaction was reduced for YFP-AtRPAP3mut2 (Figure 6D). These results confirm that the contacts of AtRPAP3 R428 and M431 with AtRuvBL2a, which we identified in the cryoEM analysis, are important for the interaction of the two proteins *in planta*.

**Figure 7.**
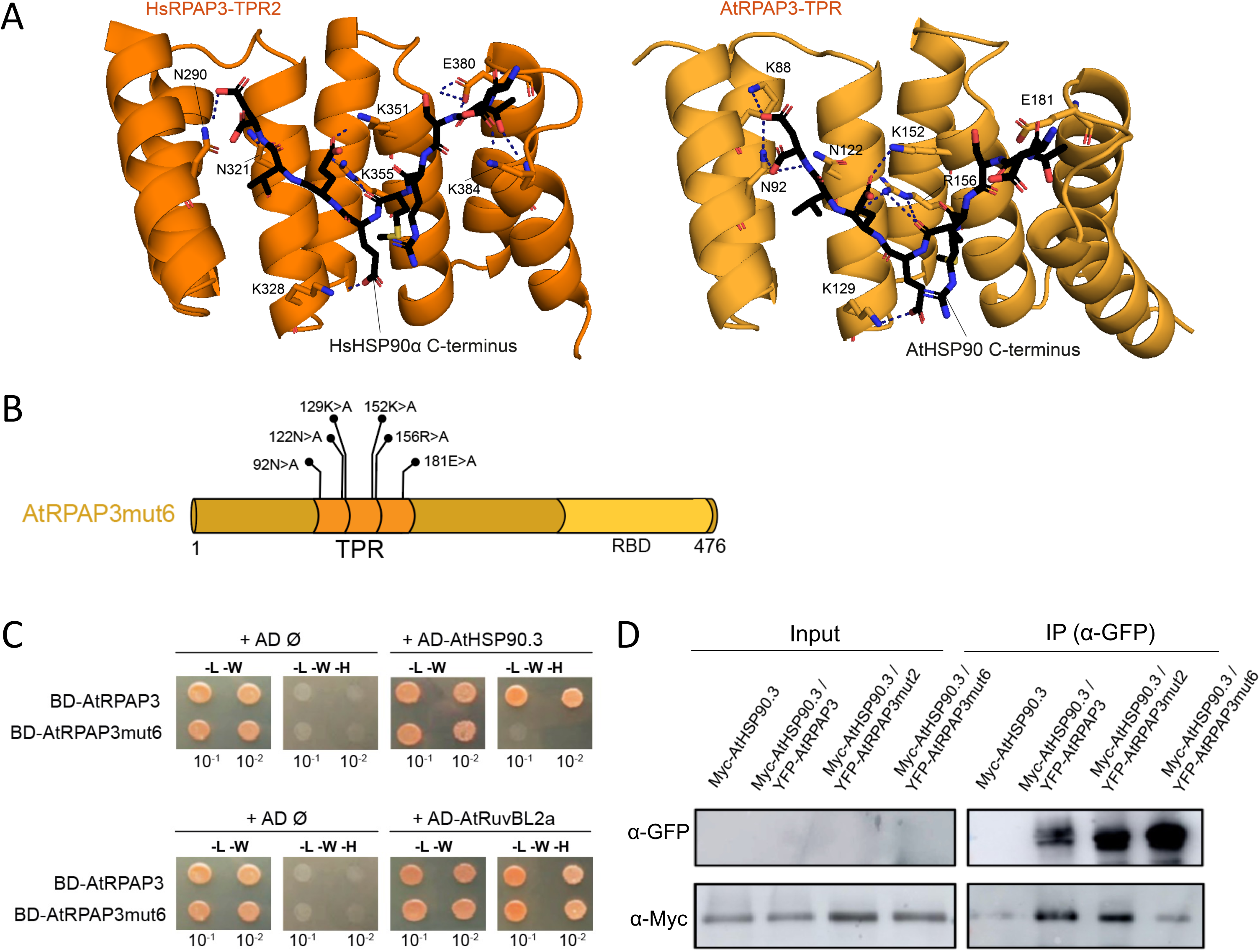
The TPR domain of AtRPAP3 mediates interaction with HSP90. (A) Close-up views of the second TPR domain of human RPAP3 and the polar contacts with the C-terminus of HsHSP90α (left) and the TPR domain of AtRPAP3 and the predicted polar contacts with the C-terminus of AtHSP90 (right). (B) Schematic of AtRPAP3mut6, a mutant version of AtRPAP3, indicating the amino acid changes in the TPR domain. (C) Y2H assays for the interaction between AtRPAP3 and AtRPAP3mut6, both fused to the GAL4 DNA-binding domain (BD), and AtRuvBL2a and AtHSP90.3, both fused to the GAL4 activation domain (AD). L, leucine; W, tryptophan; H, histidine. The numbers indicate the dilution used in the assay. (D) Co-immunoprecipitation assay showing the interaction between AtRPAP3, AtRPAP3mut2 and AtRPAP3mut6 fused to YFP, and AtHSP90.3 fused to Myc tag in *N. benthamiana*. Total proteins were immunoprecipitated with anti-GFP antibody-coated magnetic beads from extracts of infiltrated leaves. The proteins were detected with anti-GFP and anti-Myc antibodies.

### The TPR domain of AtRPAP3 mediates the interaction with HSP90

As shown above, AtRPAP3 interacts with AtHSP90.3 (Figures 2, 4E and 6C, and Supplementary Figure 3). Structural analyses indicate that the conserved MEEVD C-terminal peptide of HSP90 contacts human RPAP3 via the TPR2 domain (Henri et al., 2018). To determine whether the interaction mechanism of RPAP3 and HSP90 is conserved in plants, we modeled the structure of the TPR domain of AtRPAP3 bound to the MEEVD peptide based on the structure of the peptide bound to human RPAP3 TPR2 (Henri et al., 2018), as we could not detect the TPR of AtRPAP3 in the cryoEM analysis because it is located in a region with high flexibility. Modeling showed that the AtRPAP3 TPR can adopt a 3D structure similar to that of human RPAP3 TPR2 (Figure 7A), as shown for the TPR domain of sorghum RPAP3 (Machado Antonio et al., 2023). Alignment of RPAP3 sequences from humans and *Arabidopsis* showed that the six residues of human TPR2 that contact HSP90 (Henri et al., 2018) are conserved in AtRPAP3 TPR (Supplementary Figure 4; marked with red asterisks). Consistent with this, modeling showed that the MEEVD peptide also contacts the six residues in *Arabidopsis* TPR (Figure 7A).

We next determined the importance of these residues for the interaction between AtRPAP3 and AtHSP90.3 *in vivo*. For that purpose, we designed an AtRPAP3 version in which the six residues were changed to A, which we named AtRPAP3mut6 (Figure 7B). We first tested the interactions by Y2H. As shown in Figure 7C, AtRPAP3mut6 was unable to interact with AtHSP90.3, while it retained the ability to interact with AtRuvBL2a (see also Figure 6D). These results showed that mutation of the six residues impairs specifically the interaction with AtHSP90.3. Next, we tested whether the negative effect of mut6 on the interaction was also observed in plant cells. We expressed myc-AtHSP90.3 alone or together with YFP-AtRPAP3, YFP-AtRPAP3mut6 or YFP-AtRPAP3mut2 in leaves of *N. benthamiana*. We could not detect the different AtRPAP3 versions in the input lanes of the immunoblot, but they were detected after IP with anti-GFP antibodies (Figure 7D). Consistent with the results in yeast, myc-AtHSP90.3 was efficiently immunoprecipitated with anti-GFP antibodies from extracts of leaves co-expressing either YFP-AtRPAP3 or YFP-AtRPAP3mut2, but not from extracts co-expressing AtRPAP3mut6, in which the intensity of the myc-AtHSP90.3 band matched that of the negative control (Figure 7D). These results suggest that *Arabidopsis* RPAP3 interacts with HSP90 via the same mechanism as its orthologs in humans. In addition, our results showing the specific effect of the mut2 and mut6 versions of AtRPAP3 also suggest the absence of crosstalk between the RBD and TPR domain. This idea aligns with our structural data suggesting that the TPR domain of AtRPAP3 is in a flexible region that likely buffers any reciprocal effects that mutations in the RBD or TPR may exert.

### AtRPAP3 is required for the function of HSP90

We aimed next at determining the importance of AtRPAP3 for HSP90 function in *Arabidopsis*. As proxy to disturb HSP90 function we used geldanamycin (GDA), a natural compound that inhibits the binding of ATP to the catalytic pocket of the chaperone and thus blocks its conformational cycle in an open state (Roe et al., 1999; Schopf et al., 2017). GDA has been used in *Arabidopsis* to demonstrate, for example, a role for HSP90 in buffering the effects of genetic variation (Queitsch et al., 2002) or in stabilizing the LSM2-8 splicing complex (Esteve-Bruna et al., 2020). We treated two-day-old WT seedlings and seedlings with the *AtRPAP3* mutant allele *tpr5-2* (Sotta et al., 2016) with mock solution or 10 μM GDA and analyzed the morphological changes after ten days of treatment using AI-assisted object detection (Figures 8A and 8B and Supplementary Figure S10). Remarkably, the growth of the mutant seedlings was less affected by GDA than that of the WT when measuring the seedling area. This result indicates that RPAP3 activity is required for HSP90 to fully respond to GDA.

**Figure 8.**
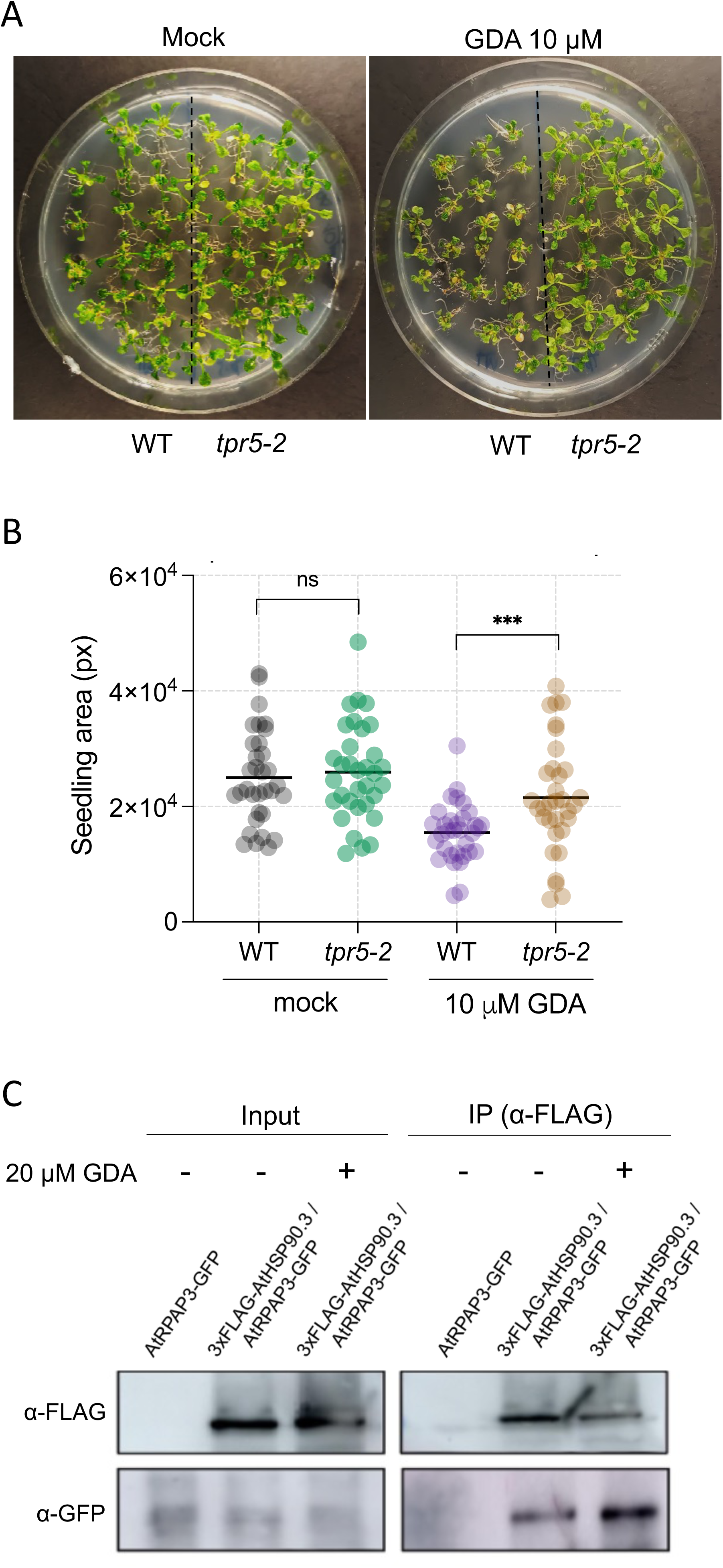
AtRPAP3 is required for the function of AtHSP90. (A) Fourteen-day-old WT and *trp5-2* seedlings treated with 10 μM GDA. (B) Total area of 14-day-old seedlings, untreated and treated with 10 μM GDA. 33 seedlings per genotype and treatment were measured. Data from one of two replicates are shown. The total area was measured with BioDock. ***, *P* < 0.001; ns, not significant. (C) Co-immunoprecipitation assay showing the interaction between AtRPAP3-GFP and 6xHis-TEV-3xFLAG-AtHSP90.3 (labeled as 3xFLAG-AtHSP90.3 in the figure) in *Arabidopsis* seedlings. Total proteins were immunoprecipitated with anti-FLAG antibody-coated magnetic beads from extracts of 7-day-old seedlings, treated with 20 µM GDA for 24 h or untreated. The proteins were detected with anti-GFP and anti-FLAG antibodies.

To gain insight into the possible molecular basis of the tolerance to GDA observed in the *tpr5* mutant, we investigated whether the inhibitor affects the interaction between AtRPAP3 and AtHSP90.3 in *Arabidopsis* seedlings. To this end, we generated and expressed a *pAtHSP90.3:6xHis-TEV-3xFLAG-AtHSP90.3* construct in the *pAtRPAP3:AtRPAP3-GFP tpr5* background (Sotta et al., 2016). We inserted the 6xHis-TEV-3xFLAG tag at the N-terminus of the chaperone because the C-terminal tag may interfere with interaction with partners, especially those that interact with the C-terminal end, such as RPAP3, as has been observed in humans (Gano and Simon, 2010). Seven-day-old seedlings expressing AtRPAP3-GFP or 6xHis-TEV-3xFLAG-AtHSP90.3/AtRPAP3-GFP fusion proteins were treated with mock solution or 20 μM GDA for 24 h and the interaction between both proteins analyzed by co-IP. Interestingly, AtRPAP3-GFP was more efficiently co-immunoprecipitated by anti-FLAG antibodies from samples treated with GDA (Figure 8C). This result is consistent with the observation that in human HEK293T cells, the interaction of HSP90 with RPAP3 and its partners PIH1D1 and URI1 is preferentially detected in cells treated with GDA (Gano and Simon, 2010). It is therefore reasonable to assume that the binding of the R2T complex to AtHSP90 via AtRPAP3 is required for the chaperone to reach a stage in its cycle that makes it sensitive to GDA. The R2T may therefore bind AtHSP90 when the chaperone is in an open conformation, like the RPAP3-related cochaperone Sti1/HOP, which binds HSP90 until another cochaperone, p23, and ATP displace it and the cycle continues to reach a closed conformation (Li et al., 2011). The R2T may contribute to HSP90 being ready to bind ATP, or eventually GDA. The reduced activity of AtRPAP3 in the *tpr5* mutant likely makes AtHSP90 less susceptible to binding of GDA, hence the tolerance shown.

## DISCUSSION

We have shown that the R2T co-chaperone complex is present in *Arabidopsis* and that it associates with the PFDL complex, as found in animals. The identification of subunits of both complexes in reciprocal immunoprecipitations, especially after TAP, suggests that both can be closely associated with each other. In fact, pairwise interaction analysis using Y2H showed that several subunits of both complexes are involved in making direct contacts. Based on interactome analyses, it has been hypothesized that the PFDLc associated with the R2TP is involved in the assembly of human RNA polymerase II, although the exact role remains unclear (Boulon et al., 2010). The association between the two complexes could also be important for the assembly of nuclear RNA polymerases in plants, as different subunits of the polymerases were immunoprecipitated by AtRPAP3, PFD6 or both (Supplementary Figure S2).

The key feature of the R2T complex found in this work is that its structure is substantially different to that of the related R2TP co-chaperone complex. R2T is organized in a double ring of AtRuvBL1-AtRuvBL2a, while the R2TP contains a single ring (Rivera-Calzada et al., 2017; Martino et al., 2018; Muñoz-Hernández et al., 2019). While the DII side of the ring in yeast and humans is occupied by Tah1-Pih1 and RPAP3-PIH1D1, respectively, which prevents dodecamerization and disrupts preformed double-ring complexes, the absence of Pih1/PIH1D1 homologs in *Arabidopsis* allows the AtRuvBL1-AtRuvBL2a complex to remain in its dodecameric form after binding to AtRPAP3. What is the functional contribution of the dodecameric versus the hexameric organization for R2T and possibly R2TP complexes? Studies in yeast have shed some light on the possible role of the two alternative organizations, as RvB1-RvB2 can switch between hexamer and dodecamer in the context of INO80-C assembly (Zhou et al., 2017). The interaction of a specific domain of the Ino80 subunit with the DII domains of Rvb1-Rvb2 triggers dodecamerization and ATPase activity, which leads to the subsequent separation of the two rings, one of them attached to Ino80. Thus, the dodecamer-hexamer cycle depends on the insertion of a client protein -Ino80-into the flexible region between the two rings, which facilitates the incorporation of the Ino80-RvB1-RvB2 subassembly into the INO80 complex. RPAP3 neither alters the ATPase activity nor disrupts the dodecamer in *Arabidopsis* (this work) nor alters ATPase activity in humans (Wing et al., 2022). However, it is the binding of Pih1/PIH1D1 that prevents the formation of the dodecamer and promotes ATPase activity in yeast (Rivera-Calzada et al., 2017) and has the potential to do so in humans (Muñoz-Hernández et al., 2019). It is possible that the alternate binding of Pih1-Tah1 to Rvb1-Rvb2 and of PIH1D1 to anchored RPAP3 triggers the alternate disruption of the dodecamer *in vivo*, promoting the ATPase cycle and facilitating the transfer of clients recruited by Pih1/PIH1D1 to HSP90 in yeast and humans. Our structural analysis shows that the two rings of *Arabidopsis* R2T are flexibly bound to each other through interactions involving the DII domains of AtRuvBL1-AtRuvBL2a. This opens up the possibility that the interaction of a client protein with the DII domains triggers the ATPase cycle and the separation of the two rings, facilitating the transfer of the client to AtHSP90. Alternatively, in the absence of a PIH1D1 homolog, AtRPAP3 could trigger the ATPase cycle and disruption of the dodecamer upon binding of client proteins. Indeed, both the RBD and the N-terminal domain of human RPAP3 are able to establish specific interactions with potential clients or adaptor proteins (Maurizy et al., 2018; Wing et al., 2022). In addition, we have shown here that various subunits of the PFDLc interact directly with AtRuvBL2a and AtRPAP3, so it is conceivable that recruitment of PFDLc contributes to the potential dodecamer-hexamer cycle.

In addition, the R2T complex is built containing only one RBD per AtRuvbl1-AtRuvBL2a ring. This is in contrast to human R2TP, where the hexameric ring harbors 1, 2 or 3 RBDs, one per RuvBL2 subunit (Martino et al., 2018; Muñoz-Hernández et al., 2019). If the binding of AtRPAP3 to each AtRuvBL2a subunit were completely independent, one would expect there to be at least a small population of molecules harboring more than one RBD per ring. However, our results are consistent with a model in which binding of the first AtRPAP3 molecule renders further binding events less favorable. Interestingly, R2T complex 1 contains 2 RBD domains, one per ring. If the interaction of AtRPAP3 with each of the rings were completely independent, one would expect that for a given position of the RBD in one of the rings, we would have found more than one possible position for the RBD in the opposite ring, which is not the case. Overall, the structure of R2T suggests the existence of allosteric communication between the AtRuvBL2a molecules within each ring and between the two rings. We cannot exclude the possibility that the different occupancy for RPAP3 between human R2TP and *Arabidopsis* R2T may be due in part to the different methodology used to reconstitute the two complexes. Human R2TP was prepared by mixing an excess of RPAP3 with RuvBL1-RuvBL2, whereas *Arabidopsis* R2T was produced by co-expressing all subunits in cells. In either case, it can be argued that this latter method may better reflect what might happen *in vivo*.

In many species, including *Arabidopsis*, the presence of the R2T complex is associated with the loss of Pih1/PIH1D1 orthologs. Phylogenetic analyses have suggested that Pih1/PIH1D1 orthologs have been lost in species that lack cilia (Maurizy et al., 2018). This suggests that the role of the R2TP in the assembly of axonemal dyneins involved in cilia formation has been a major force in maintaining the full complex during evolution (Lynham and Houry, 2022). The two Pih1/PIH1D1 homologs in the liverwort *Marchantia polymorpha* (Bowman et al., 2017) are expressed exclusively in the antheridium, the male sex organ that produces ciliated sperm cells, while expression of the homologous gene of *RPAP3* is broader (Supplementary Figure S11), consistent with the idea of a specialized function of the R2TP complex in plants, with the R2T fulfilling all other functions. Although the phylogenetic analysis of Maurizy et al. (2018) suggests that only bryophytes have Pih1/PIH1D1 orthologs within the green lineage, orthologs can be found in non-seed tracheophytes such as the lycophyte *Selaginella moellendorffii* (Banks et al., 2011) and the fern *Ceraptopteris richardii* (Marchant et al., 2021), both of which have cilia, e.g. in sperm cells (Hodges et al., 2012). The specialization of the R2TP complex in non-seed plants may have been caused by the transcriptional restriction of *Pih1/PIH1D1* homolog genes in the male reproductive organs. Both R2T and R2TP complexes could coexist in these organs if the R2T can be formed by competition between RPAP3 and Pih1/PIH1D1 homologs by binding to RuvBL1-RuvBL2 as in humans (Wing et al., 2022). The specialized function of the R2TP complex in non-seed plants and its absence in seed plants such as *Arabidopsis* suggests that the clients normally recruited by Pih1/PIH1D1 either have no orthologs in plants or their function does not require the involvement of the complex. Overall, this may have led to an expansion of the function of the R2T in plants to fulfill the basic functions of the R2TP. In humans, the R2T exerts its function mainly in the nucleus, as the RPAP3-2 isoform is a nuclear protein, whereas the full-length RPAP3 is mainly cytoplasmic (Maurizy et al., 2018). We have shown here that AtRPAP3 is localized in both the nucleus and the cytoplasm. This suggests that a change in the subcellular localization of plant RPAP3 may have originally been associated with the specialization of the R2TP in non-seed plants, which has been retained after the loss of Pih1/PIH1D1 in seed plants.

## METHODS

### Plant material, growth conditions and treatments

*Arabidopsis* mutants and transgenic lines are in Col-0 background. The *AtRPAP3* mutant allele *tpr5-2* and the transgenic line *pAtRPAP3:AtRPAP3-GFP tpr5-2* have been described (Sotta et al., 2016). We prepared an *Arabidopsis* line expressing *pAtHSP90.3:6xHis-TEV-3xFLAG-AtHSP90.3* in the *pAtRPAP3:AtRPAP3-GFP tpr5-2* background. To make the *pAtHSP90.3:6xHis-TEV-3xFLAG-AtHSP90.3* construct, we amplified by PCR a genomic fragment starting -1141 bp upstream of the *AtHSP90.3* transcription start site until the stop codon, which was included, and was cloned by Gateway first into the pDONR207 and then into the destination vector pMDC123 (Curtis and Grossniklaus, 2003). Next, a NcoI site was introduced at the starting ATG by PCR, which was used to insert a DNA fragment containing the *6xHis-TEV-3xFLAG* sequence by In-Fusion technology. This DNA fragment encodes the TEV protease recognition site between the 6xHis and 3xFLAG sequences. The construct was introduced into *pAtRPAP3:AtRPAP3-GFP tpr5-2* plants via *Agrobacterium tumefaciens* strain C58 transformation. Transgenic plants were selected in 1/2 Murashige and Skoog (MS) medium containing basta herbicide. All oligonucleotides used for cloning are shown in the table in Supplementary Table S2.

### Identification of PFD6 and AtRPAP3 interactors

The CDS with stop codon of AtRPAP3 was amplified by PCR from a cDNA pool of one-week-old *Arabidopsis* seedlings, cloned into pDONR207 and transferred into the pKNGS_rhino vector (Van Leene et al., 2015) by Gateway to create a GS-AtRPAP3 fusion. The GS-PFD6 construct was prepared using pENTR223-PFD6 (Esteve-Bruna et al., 2020). As control construct, pKGNS_rhino fused to a nonsense stuffer sequence was used to express GS alone. *Agrobacterium tumefaciens* C58 cells carrying pKNGS_rhino-PFD6, pKNGS_rhino-AtRPAP3 and the control construct were used to transform PSB-D cell suspensions (Van Leene et al., 2015). 11 g of cells expressing pKNGS_rhino-PFD6 from each of two biological replicates were used for tandem affinity purification (Van Leene et al., 2015). Single immunoprecipitations (IP) with 7 g of cells expressing pKNGS_rhino-AtRPAP3 and the control construct from each of the three biological replicates were performed according to the previously described protocol (Antosz et al., 2017). Interactors of GS-PFD6 were identified by mass spectrometry at the Proteomics Facility of the Centro Nacional de Biotecnología (Madrid, Spain). Proteins identified as GS-PFD6 interactors were those not included in the list of background proteins that appear non-specific in TAP experiments with the same tag (Van Leene et al., 2015). Proteins pulled down by GS-AtRPAP3 and GS were identified by mass spectrometry at the Servicio de Proteómica of the Universidad de Córdoba (Spain). The proteins identified as GS-AtRPAP3 interactors were those whose peptides were enriched at least 5 times compared to the peptides found in the control GS IP.

### Yeast two-hybrid assays

The CDSs with stop codon of AtRuvBL1, AtRuvBL2a, AtHSP90-3 and ASDURF were amplified by PCR from a cDNA pool of one-week-old *Arabidopsis* seedlings and cloned by Gateway into pDONR207. The AtRPAP3mut2 version, in which the codons encoding R428 and M431 were changed to encode A, was produced by overlapping PCR and cloned into pDONR207. The CDS of the AtRPAP3mut6 version, in which the codons encoding N92, N122, K129, K152, R156 and E181 were changed to encode A, was synthesized by Integrated DNA Technologies (Belgium) and cloned into pDONR207. The CDSs of AtRPAP3, AtRuvBL1, AtRuvBL2a, AtHSP90-3 and ASDURF were transferred to the pGADT7-GW and pGBKT7-GW destination vectors using Gateway LR Clonase II enzyme mix to produce fusion proteins of the Gal4-activation domain (AD) and GAL4 DNA-binding domain (BD), respectively. The CDS of AtURI, UXT, PFD2 and PFD6 cloned into pGADT7-GW and pGBKT7-GW have been previously described (Gómez-Mínguez et al., 2024). The CDSs of AtRPAP3mut2 and AtRPAP3mut6 were transferred to the pGBKT7-GW destination vector using Gateway LR Clonase II enzyme mix. The expression vectors derived from pGADT7 and pGBK7 were introduced into Y187 and Y2HGold yeast strains, respectively. The transformants were selected with Synthetic Defined (SD) medium, which lacked leucine (-Leu) or tryptophan (-Trp). Haploid yeast cells were mated to obtain diploid cells by selection in SD-Leu-Trp medium. Protein interactions were tested by the nutrient requirement of histidine (His) in SD-Leu-Trp-His plates.

### Confocal microscopy

The CDSs of AtRPAP3, AtRuvBL2a and AtHSP90.3 were transferred by Gateway into the pEarleyGate-104 vector (Earley et al., 2006) to fuse YFP to the N-terminal end of each protein. The AtRPAP3 CDS was also transferred into the vector pH7WGR2 (gatewayvectors.vib.be/collection/ph7wgr2) to fuse mRFP to its N-terminus. Leaves of three-week-old *Nicotiana benthamiana* plants grown under a 16 h light: 8 h dark photoperiod were infiltrated with *Agrobacterium tumefaciens* C58 carrying constructs to express AtRPAP3, AtRuvBL2a or AtHSP90-3 fused to YFP or RFP, and the p19 silencing suppressor. Leaf confocal fluorescence was recorded on the third day after agroinfiltration using a Zeiss Axio-Observer 780 Pascal confocal microscope. The YFP signal was excited with an argon laser at the wavelength of 488 nm, and its emission was detected at 505-530 nm. The RFP signal was excited with an argon laser at the wavelength of 555 nm, and its emission was detected at 575-600 nm. Chloroplasts autofluorescence was detected between 675 and 760 nm.

### Bimolecular fluorescence complementation assay

The CDS of AtRPAP3 and that of AtRuvBL2a and AtHSP90.3 were transferred into pMDC43-YFN and pMDC43-YFC vectors, respectively (Belda-Palazón et al., 2012), by Gateway for bimolecular fluorescence complementation (BiFC) assays. *Agrobacterium tumefaciens* C58 cultures containing combinations of these plasmids and those of negative controls (at a ratio of 1:1 up to a total optical density of 0.1) were used to infiltrate 3-week-old *N. benthamiana* leaves. Three days after infiltration, confocal images were collected using Zeiss Axio-Observer 780 Pascal confocal microscope. Reconstituted YFP signal was detected in abaxial epidermal cells at 503-517 nm after excitation with an Argon laser (488 nm).

### Cloning, expression and purification of recombinant proteins

The CDS of AtRuvBL1 was cloned into a modified pET-15b vector (Novagen) including a N-terminal histidine tag followed by tobacco etch virus (TEV) protease site, and untagged AtRuvBL2a was cloned into pCDFDuet-1 vector (Novagen) using standard cloning protocols. AtRPAP3 was cloned into pRSFDuet-1 vector (Novagen) with a N-terminal His-tag followed by an IF2D1 tag (translation initiation factor 2 domain 1) and a TEV protease site, and a C-terminal Strep-tag with human rhinovirus 3C (HRV 3C) protease site using the IVA cloning methodology (García-Nafría et al., 2016).

Expression of the AtRuvBL1-AtRuvBL2a complex was carried out by co-expression of the two plasmids containing the individual proteins in *E. coli* LOBSTR (DE3) cells and induction with 0.1 mM isopropyl-β-D-1-thiogalactopyranoside (IPTG) for 4 h at 28°C. Cells were collected by centrifugation at 5,000 rpm and 4°C, and lysed in 50 mM Tris-HCl, 300 mM NaCl, 10% (v/v) glycerol, 0.1% (v/v) NP-40 buffer supplemented with lysozyme (0.3 mg/ml), benzonase (250 U/µl) (Novagen) and cOmplete^®^ EDTA-free protease Inhibitor Cocktail (Roche) by sonication. After clarification at 50,000 rpm during 1 h at 4°C, lysate was filtered through a 0.45 µm device and loaded onto a HisTrap HP (Cytiva) column equilibrated in 40 mM Tris-HCl, 200 mM NaCl, 5% (v/v) glycerol, 20 mM imidazole buffer. Elution was performed using a 20-500 mM imidazole gradient, and fractions containing the AtRuvBL1-AtRuvBL2a complex were pooled and dialyzed in 50 mM Tris-HCl pH 7.4, 150 mM NaCl, 10% (v/v) glycerol buffer over night at 4°C. The sample was further purified using a Superose 6 Increase GL 3.2/100 column (Cytiva) equilibrated in 50 mM Tris-HCl pH 7.4, 150 mM NaCl buffer.

AtRPAP3 was produced by expression in *E. coli* BL21 (DE3) strain by induction with 0.25 mM IPTG at 16°C for 18 h. Cells were harvested and lysed by sonication in 50 mM HEPES, 500 mM NaCl, 10% (v/v) glycerol, 0.5% (v/v) NP-40, 1 mM DTT buffer supplemented with lysozyme (0.3 mg/ml), benzonase (250 U/µl) (Novagen) and cOmplete^®^ EDTA-free protease Inhibitor Cocktail (Roche). Cleared lysate was loaded into a HisTrap HP (Cytiva) column equilibrated in 25 mM HEPES pH 8, 300 mM NaCl, 20 mM imidazole, and purified protein was eluted with a 20-500 mM imidazole gradient. Pooled fractions containing AtRPAP3 were loaded onto a StrepTactin^®^XT 4Flow^®^ high-capacity column (IBA Lifesciences) equilibrated in 25 mM HEPES pH 8, 300 mM NaCl, 1 mM DTT buffer and eluted with the same buffer supplemented with 50 mM biotin. Purified AtRPAP3 was concentrated with an Amicon^®^ Ultra-4 30,000 MWCO centrifugal filter unit (Millipore) and size exclusion purified in a Superose 6 Increase 10/300 GL (Cytiva) column equilibrated in 25 mM HEPES pH 8, 300 mM NaCl, 1 mM DTT.

The AtRuvBL1-AtRuvBL2a-AtRPAP3 (R2T) complex was produced by co-expression of AtRuvBL1, AtRuvBL2a and AtRPAP3 in *E. coli* BL21 (DE3) cells and induction with 0.25 mM IPTG at 17°C for 18 h. The complex was purified by pulling down from the Strep-tag in AtRPAP3 with a StrepTactin^®^XT 4Flow^®^ column (IBA Lifesciences) using the same protocol described for AtRPAP3. Fractions containing the R2T complex were dialyzed in 25 mM HEPES pH 8, 300 mM NaCl, 1 mM DTT, 10% (v/v) glycerol buffer, and further purified in a Superose 6 Increase 10/300 GL (Cytiva) column equilibrated in 25 mM HEPES pH 8, 300 mM NaCl, 1 mM DTT.

### ATPase assays

Activity of the purified AtRuvBL1-AtRuvBL2a complex was tested by measuring the ATP hydrolysis in a continuous spectrophotometric pyruvate kinase-lactate dehydrogenase-coupled assay, based on the regeneration of the hydrolyzed ATP coupled to oxidation of NADH. NADH absorbance at 340 nm was measured using a CLARIOstar Plus plate reader (BMG Labtech) in time course experiments, and ATP hydrolysis rates were calculated from the slope of the decreasing curves. Assays were performed at 37°C in 100 μl reactions in buffer 50 mM Tris-HCl pH 7.4, 150 mM NaCl, 20 mM MgCl_2_, containing 2 mM phosphoenolpyruvate (PEP), 0.5 mM NADH, 0.04 U/μl pyruvate kinase/0.05 U/μl lactic dehydrogenase (Sigma-Aldrich) and 5 mM ATP. Hydrolysis reactions were started by addition of 3 μM of AtRuvBL1-AtRuvBL2a (concentration calculated considering monomers), 1 μM AtRPAP3 (control experiment) or 3 μM of AtRuvBL1-AtRuvBL2a plus 1 μM AtRPAP3, and carried out for 50 min. ATP turnover (mol ATP/mol protein) indicated in min^−1^ was calculated for a time interval during which the absorbance decrease was linear. Values in the graph are indicated as percentage of the rate of the wild-type protein.

### CryoEM

3 μl of freshly purified protein complex, either AtRuvBL1-AtRuvBL2a or R2T, at 0.8 mg/ml were applied to glow discharged Quantifoil^®^ 300 mesh R 0.6/1 grids and flash frozen in liquid ethane using FEI Vitrobot MAG IV (Thermo Fisher Scientific). For AtRuvBL1-AtRuvBL2a complex, 1001 movies were collected on a JEOL JEM-2200FS microscope operating at 200 kV, equipped with a K3 counting camera (Gatan). Calibration and automatic data collection were performed with SerialEM (University of Colorado) at a nominal magnification of ×40k, physical pixel size of 0.9772 Å, a total dose of 40 e^−^/Å^2^, and 40 fractions. Autofocus was performed using an objective defocus range between −1.5 and −3.0 μm (Supplementary Table S1).

For the R2T complex, 6139 movies were collected in a Titan Krios equipped with a Falcon 4i detector (Thermo Fisher Scientific) and Selectris X energy filter at eBIC (Diamond Light Source, Oxford, UK) in counting mode, and a slit width of 5 eV. Microscope calibrations and automatic data acquisition were performed with EPU software (Thermo Fisher Scientific) at a nominal magnification of ×130k, physical pixel size of 0.921 Å, a total dose of 50.12 e^−^/Å^2^, 8.05 e^−^/Å^2^/s and 50 fractions. Autofocus was performed using an objective defocus range between −1.5 and −3.0 μm. CryoEM images were collected at the Diamond Light Source cryoEM facility at the UK’s National Electron Bio-imaging Center (eBIC).

### Image processing

Local drift was corrected on images of the AtRuvBL1-AtRuvBL2a complex using MotionCor2 (5 × 5 patches) and dose-weighting of fraction stacks (Zheng et al., 2017). Contrast transfer function (CTF) parameters of drift-corrected images were determined with Gctf (Zhang, 2016). The 179,410 best quality particles were selected after 2D classification in RELION of the initial data set containing 297,450 particles (Supplementary Figure S5C). Initial model was obtained using cryoSPARC (Punjani et al., 2017), and 3D refined with RELION. After 3D classification, the best particles were selected and further refined. Focus 3D refinement on the top and bottom rings of the dodecameric volume was performed to obtain 4.7 Å and 6.7 Å maps, respectively, using gold-standard methods and the Fourier Shell Correlation (FSC) cut-off of 0.143.

Images of the R2T complex were local drift corrected and CTF parameters were determined as described. A subset of manually picked particles was used to generate reference-free 2D averages in RELION (Zivanov et al., 2018) that were further used for template-based automatic particle picking. R2T complex initial data set containing 2,334,954 particles was 2D-classified in RELION, and the 788,672 best quality particles were selected (Supplementary Figure S8, Supplementary Table S1). *Ab initio* 3D model was generated from the selected particles using cryoSPARC (Punjani et al., 2017), and used as reference for a consensus 3D-refinement in RELION. A subset of 481,879 particles was selected after getting rid of junk particles by 3D-classification of the dodecameric R2T model and further 3D-refined. Densities corresponding to the RBD domain of AtRPAP3 bound to the AtRuvBL1-AtRuvBL2a dodecamer were poorly defined, so a focus 3D-classification protocol was applied using a mask containing the hexameric top ring and three RBD densities, defining a population of particles with no AtRPAP3 bound (296,837 particles, 61.6 %) and 3 populations containing one RBD molecule bound each in different positions (185,042 particles, 38.4 %) (Supplementary Figure S8). The RBD-containing particles were aligned to the same position based on the RBD density and focus 3D-refined. After applying the same focus 3D-classification protocol to the bottom hexamer of the complex, two dodecameric populations were identified, one containing a second RBD density in the bottom hexamer (complex 1, 7.75 Å; 18%) and one with no density for RBD in bottom ring (complex 2, 8.18 Å; 82%). Particles with the top ring RBD density aligned were 3D-refined with a mask engaging the top ring AtR2T, and after iterative rounds of CTF refinement and Bayesian Polishing in RELION, a cryoEM map for the top ring R2T complex was obtained at an average resolution of 3.73 Å map using gold-standard methods and the Fourier Shell Correlation (FSC) cut-off of 0.143 (Supplementary Figure S9A).

### Model building

The high-resolution cryoEM map of the top ring of the R2T complex was used for model building using as starting models the atomic models of the AtRuvBL1 (AF-Q9FMR9-F1), AtRuvBL2a (AF-Q9FJW0-F1) and AtRPAP3 (AF-Q5XF05-FA) proteins generated in AlphaFold2 (Yang et al., 2023). Using the model of the human R2TP complex as template (PDB 6FO1), 3 copies of the AtRuvBL1 model, 3 copies of AtRuvBL2a and one AtAtRPAP3 RBD domain were aligned to create the hexameric complex and rigid fitted in the cryoEM map of R2T using USCF Chimera. An initial step of automatic refinement was done in phenix.real_space_refinement (Afonine et al., 2018), followed by interactive model building using ISOLDE (Croll, 2018) and manual refinement of the seven chains in Coot (Emsley et al., 2010). A final step of automatic refinement was done in phenix.real_space_refinement to improve the geometries of the model (Supplementary Table S1).

Using the TPR2 domain of human RPAP3 as a template (PDB code: 6FDP), the structure of the TPR domain of AtRPAP3 was generated using Modeler (version 9.23) (Webb and Sali, 2016). The predicted structure of the TPR domain of AtRPAP3 was aligned with the human model (PDB code: 6FDP) and polar contacts between the TPR domain of AtRPAP3 and the MEEVD peptide were identified using PyMOL 2.4 software (pymol.org).

### Protein co-immunoprecipitation assays

The CDSs of AtRuvBL2a and AtHSP90.3 were transferred into the pEarleyGate-203 vector to create Myc fusions (Earley et al., 2006). The CDSs of AtRPAP3mut2 and AtRPAP3mut6 were transferred into pEarleyGate-104 to produce the YFP-AtRPAP3mut2 and YFP-AtRPAP3mut6 fusions by Gateway.

Leaves of three-week-old *N. benthamiana* plants were infiltrated with different mixtures of *Agrobacterium tumefaciens* C58 cultures carrying the expression vectors and the p19 silencing suppressor. Three days after infiltration, the leaves were collected and frozen in liquid nitrogen. 1 ml of the frozen tissue was homogenized in 0.5 ml of extraction buffer containing 50 mM Tris-HCl pH 7.5, 150 mM NaCl, 0.25% (v/v) Triton X-100, 2 mM PMSF and 1X cOmplete® EDTA-free protease inhibitor cocktail (Roche). The extracts were placed on ice for 15 minutes and then centrifuged 2 times at maximum speed in a benchtop centrifuge at 4°C. 150 μg of total proteins were denatured in 1 volume of 2X Laemmli buffer and kept for later use as input samples. In addition, 1 mg of total protein was incubated with 25 μl of GFP-Trap® Magnetic Beads (ChromoTek) in 1 ml of extraction buffer at 4 °C for 2 h in a rotating wheel. The beads were separated with a magnet and the supernatant was discarded. The beads were resuspended in 1 ml wash buffer (Tris-HCl pH 7.5, 350 mM NaCl, 0.25% Triton X-100 [v/v]) and incubated at 4°C for 5 min in a rotating wheel. This washing step was repeated four times. The proteins were then eluted by incubating the beads with 70 μl 2X Laemmli for 5 minutes at 95 °C. For analysis, 63 μl of the immunoprecipitated samples (representing 90% of the total) were loaded onto a 10% SDS-PAGE together with 50 μg of the input sample. After electrophoresis, the proteins were transferred to a PVDF membrane and immunodetected with an anti-Myc-HRP antibody (Myc-HRP, 1:5000, Roche). The remaining 10% of the immunoprecipitated samples together with 50 μg of the input were subjected to the same procedure but probed with an anti-GFP antibody (JL-8, 1:5000, Living Colors). After incubation with SuperSignal^TM^ West Femto (Thermo-Fisher Scientific), chemiluminescence was visualized with the ImageQuant 800 (Amersham).

To test the interaction between AtRPAP3 and AtHSP90.3 in the presence or absence of GDA, *Arabidopsis* lines expressing AtRPAP3-GFP/6xHis-TEV-3xFLAG-AtHSP90.3 and AtRPAP3-GFP (control) were used. 7-day-old *Arabidopsis* seedlings grown in 1/2 MS plates under continuous light (50–60 μmol m^−2^ s^−1^) at 22°C were transferred to liquid 1/2 MS medium containing 20 µM GDA or mock solution (DMSO) for 24 h in darkness. Total proteins were immunoprecipitated with anti-FLAG antibody-coated magnetic beads. The proteins were detected with anti-GFP and anti-FLAG antibodies. The procedure was performed as described above.

### Growth assay in response to geldanamycin

Seeds of WT and *tpr5-2* were germinated for 2 days under continuous light (50–60 μmol m^−2^ s^−1^) at 22°C. After two days, seedlings were transferred to plates containing 1/2 MS medium supplemented with a mock solution (DMSO) or 10 μM GDA (MedChem, HY-15230) and maintained under the same conditions. Photos were taken 10 days after treatment. Biodock for AI image analysis (biodock.ai) was used to quantify the area of each plant for each genotype and condition. A Tukey’s Multiple Comparison Test (One-way ANOVA) was performed and area values were plotted using Graphpad Prism 8.0.2.

## FUNDING

Research was funded by grants PID2019-109925GB-I00 and PID2020-114429RB-I00 from MCIN/AEI/10.13039/501100011033 to D.A and O.L., respectively. A.P-A. was supported by a Ministerio de Educación predoctoral contract (FPU17/05186). O.L. laboratory also had the support from the National Institute of Health Carlos III to CNIO. CryoEM data used in this work for the structure of R2T was obtained at the Diamond Light Source cryo-EM facility at the UK’s National Electron Bio-imaging Center (eBIC) under BAG Proposal No BI26876 "Stop cancer - structural studies of macromolecular complexes involved in cancer by cryo-EM".

## DATA AVAILABILITY

CryoEM map and the model of the R2T complex were deposited in the EMDB and PDB with accession codes EMD-19894 and PDB ID 9EQ2.

## ACKNOWLEDGEMENTS

The authors thank Prof. Toru Fujiwara (The University of Tokyo, Tokyo, Japan) for providing seeds of *tpr5-2* and *pAtRPAP3:AtRPAP3-GFP tpr5-2 Arabidopsis* lines.

## Notes

### Competing Interest Statement

The authors have declared no competing interest.

## REFERENCES

Afonine, P. V., Poon, B. K., Read, R. J., Sobolev, O. V., Terwilliger, T. C., Urzhumtsev, A., and Adams, P. D. (2018). Real-space refinement in PHENIX for cryo-EM and crystallography. Acta Crystallogr D Struct Biol 74:531–544.

Antosz, W., Pfab, A., Ehrnsberger, H. F., Holzinger, P., Kollen, K., Mortensen, S. A., Bruckmann, A., Schubert, T., Langst, G., Griesenbeck, J., et al. (2017). The Composition of the Arabidopsis RNA Polymerase II Transcript Elongation Complex Reveals the Interplay between Elongation and mRNA Processing Factors. Plant Cell 29:854–870.

Banks, J. A., Nishiyama, T., Hasebe, M., Bowman, J. L., Gribskov, M., DePamphilis, C., Albert, V. A., Aono, N., Aoyama, T., Ambrose, B. A., et al. (2011). The selaginella genome identifies genetic changes associated with the evolution of vascular plants. Science 332:960–963.

Belda-Palazón, B., Ruíz, L., Martí, E., Tárraga, S., Tiburcio, A. F., Culiáñez, F., Farràs, R., Carrasco, P., and Ferrando, A. (2012). Aminopropyltransferases involved in polyamine biosynthesis localize preferentially in the nucleus of plant cells. PLoS One 7:e46907.

Blanco-Touriñán, N., Esteve-Bruna, D., Serrano-Mislata, A., Esquinas-Ariza, R. M., Resentini, F., Forment, J., Carrasco-López, C., Novella-Rausell, C., Palacios-Abella, A., Carrasco, P., et al. (2021). A genetic approach reveals different modes of action of prefoldins. Plant Physiol 187:1534–1550.

Boulon, S., Pradet-Balade, B., Verheggen, C., Molle, D., Boireau, S., Georgieva, M., Azzag, K., Robert, M., Ahmad, Y., Neel, H., et al. (2010). HSP90 and its R2TP/Prefoldin-like cochaperone are involved in the cytoplasmic assembly of RNA polymerase II. Mol Cell 39:912–924.

Bowman, J. L., Kohchi, T., Yamato, K. T., Jenkins, J., Shu, S., Ishizaki, K., Yamaoka, S., Nishihama, R., Nakamura, Y., Berger, F., et al. (2017). Insights into Land Plant Evolution Garnered from the Marchantia polymorpha Genome. Cell 171:287–304 e15.

Candela-Ferré, J., Diego-Martín, B., Pérez-Alemany, J., and Gallego-Bartolomé, J. (2024). Mind The Gap: Epigenetic Regulation Of Chromatin Accessibility In Plants. Plant Physiol 10.1093/plphys/kiae024.

Cloutier, P., Al-Khoury, R., Lavallee-Adam, M., Faubert, D., Jiang, H., Poitras, C., Bouchard, A., Forget, D., Blanchette, M., Coulombe, B., et al. (2009). High-resolution mapping of the protein interaction network for the human transcription machinery and affinity purification of RNA polymerase II-associated complexes. Methods 48:381–386.

Cloutier, P., Poitras, C., Durand, M., Hekmat, O., Fiola-Masson, E., Bouchard, A., Faubert, D., Chabot, B., Coulombe, B., Fiola-Masson, É., et al. (2017). R2TP/Prefoldin-like component RUVBL1/RUVBL2 directly interacts with ZNHIT2 to regulate assembly of U5 small nuclear ribonucleoprotein. Nat Commun 8:15615.

Cloutier, P., Poitras, C., Faubert, D., Bouchard, A., Blanchette, M., Gauthier, M. S., and Coulombe, B. (2020). Upstream ORF-Encoded ASDURF Is a Novel Prefoldin-like Subunit of the PAQosome. J Proteome Res 19:18–27.

Croll, T. I. (2018). ISOLDE: A physically realistic environment for model building into low-resolution electron-density maps. Acta Crystallogr D Struct Biol 74:519–530.

Curtis, M. D., and Grossniklaus, U. (2003). A gateway cloning vector set for high-throughput functional analysis of genes in planta. Plant Physiol 133:462–469.

Dauden, M. I., López-Perrote, A., and Llorca, O. (2021). RUVBL1–RUVBL2 AAA-ATPase: a versatile scaffold for multiple complexes and functions. Curr Opin Struct Biol 67:78–85.

Earley, K. W., Haag, J. R., Pontes, O., Opper, K., Juehne, T., Song, K., and Pikaard, C. S. (2006). Gateway-compatible vectors for plant functional genomics and proteomics. Plant J 45:616–629.

Emsley, P., Lohkamp, B., Scott, W. G., and Cowtan, K. (2010). Features and development of Coot. Acta Crystallogr D Biol Crystallogr 66:486–501.

Esteve-Bruna, D., Carrasco-López, C., Blanco-Touriñán, N., Iserte, J., Calleja-Cabrera, J., Perea-Resa, C., Úrbez, C., Carrasco, P., Yanovsky, M. J., Blázquez, M. A., et al. (2020). Prefoldins contribute to maintaining the levels of the spliceosome LSM2-8 complex through Hsp90 in Arabidopsis. Nucleic Acids Res 48:6280–6293.

Gano, J. J., and Simon, J. A. (2010). A proteomic investigation of ligand-dependent HSP90 complexes reveals CHORDC1 as a novel ADP-dependent HSP90-interacting protein. Mol Cell Proteomics 9:255–270.

García-Nafría, J., Watson, J. F., and Greger, I. H. (2016). IVA cloning: A single-tube universal cloning system exploiting bacterial In Vivo Assembly. Sci Rep 6:27459.

Gestaut, D., Roh, S. H., Ma, B., Pintilie, G., Joachimiak, L. A., Leitner, A., Walzthoeni, T., Aebersold, R., Chiu, W., and Frydman, J. (2019). The Chaperonin TRiC/CCT Associates with Prefoldin through a Conserved Electrostatic Interface Essential for Cellular Proteostasis. Cell 177:751–765 e15.

Gómez-Mínguez, Y., Palacios-Abella, A., Costigliolo-Rojas, C., Barber, M., Hernández-Villa, L., Úrbez, C., and Alabadí, D. (2024). The prefoldin-like protein AtURI exhibits characteristics of intrinsically disordered proteins. FEBS Lett 598:556–570.

Gorynia, S., Bandeiras, T. M., Pinho, F. G., McVey, C. E., Vonrhein, C., Round, A., Svergun, D. I., Donner, P., Matias, P. M., and Carrondo, M. A. (2011). Structural and functional insights into a dodecameric molecular machine - The RuvBL1/RuvBL2 complex. J Struct Biol 176:279–291.

Gstaiger, M., Luke, B., Hess, D., Oakeley, E. J., Wirbelauer, C., Blondel, M., Vigneron, M., Peter, M., and Krek, W. (2003). Control of nutrient-sensitive transcription programs by the unconventional prefoldin URI. Science 302:1208–1212.

Henri, J., Chagot, M. E., Bourguet, M., Abel, Y., Terral, G., Maurizy, C., Aigueperse, C., Georgescauld, F., Vandermoere, F., Saint-Fort, R., et al. (2018). Deep Structural Analysis of RPAP3 and PIH1D1, Two Components of the HSP90 Co-chaperone R2TP Complex. Structure 26:1196–1209.

Hodges, M. E., Wickstead, B., Gull, K., and Langdale, J. A. (2012). The evolution of land plant cilia. New Phytologist 195:526–540.

Holt Iii, B. F., Boyes, D. C., Ellerströ, M., Siefers, N., Wiig, A., Kauffman, S., Grant, M. R., and Dangl, J. L. (2002). An Evolutionarily Conserved Mediator of Plant Disease Resistance Gene Function Is Required for Normal Arabidopsis Development. Dev Cell 2:807–817.

Horejsi, Z., Stach, L., Flower, T. G., Joshi, D., Flynn, H., Skehel, J. M., O’Reilly, N. J., Ogrodowicz, R. W., Smerdon, S. J., and Boulton, S. J. (2014). Phosphorylation-dependent PIH1D1 interactions define substrate specificity of the R2TP cochaperone complex. Cell Rep 7:19–26.

Houry, W. A., Bertrand, E., and Coulombe, B. (2018). The PAQosome, an R2TP-based chaperone for quaternary structure formation. Trends Biochem Sci 43:4–9.

Hubert, D. A., He, Y., Mcnulty, B. C., Tornero, P., and Dangl, J. L. (2009). Specific Arabidopsis HSP90.2 alleles recapitulate RAR1 cochaperone function in plant NB-LRR disease resistance protein regulation. PNAS 106:9556–9563.

Jeronimo, C., Forget, D., Bouchard, A., Li, Q., Chua, G., Poitras, C., Therien, C., Bergeron, D., Bourassa, S., Greenblatt, J., et al. (2007). Systematic analysis of the protein interaction network for the human transcription machinery reveals the identity of the 7SK capping enzyme. Mol Cell 27:262–274.

Kanemaki, M., Makino, Y., Yoshida, T., Kishimoto, T., Koga, A., Yamamoto, K., Yamamoto, M., Moncollin, V., Egly, J.-M., Muramatsu, M., et al. (1997). Molecular Cloning of a Rat 49-kDa TBP-Interacting Protein (TIP49) That Is Highly Homologous to the Bacterial RuvB. Biochem Biophys Res Commun 235:64–68.

Kim, T. S., Kim, W. Y., Fujiwara, S., Kim, J., Cha, J. Y., Park, J. H., Lee, S. Y., and Somers, D. E. (2011). HSP90 functions in the circadian clock through stabilization of the client F-box protein ZEITLUPE. Proc Natl Acad Sci U S A 108:16843–16848.

Krishna, P., and Gloor, G. (2001). The Hsp90 family of proteins in Arabidopsis thaliana. Cell Stress Chaperones 6:238–246.

Li, J., Richter, K., and Buchner, J. (2011). Mixed Hsp90-cochaperone complexes are important for the progression of the reaction cycle. Nat Struct Mol Biol 18:61–67.

López-Perrote, A., Hug, N., González-Corpas, A., Rodríguez, C. F., Serna, M., García-Martín, C., Boskovic, J., Fernandez-Leiro, R., Caceres, J. F., and Llorca, O. (2020). Regulation of RUVBL1-RUVBL2 AAA-ATPases by the nonsense-mediated mRNA decay factor DHX34, as evidenced by Cryo-EM. Elife 9:e63049.

Lynham, J., and Houry, W. A. (2022). The Role of Hsp90-R2TP in Macromolecular Complex Assembly and Stabilization. Biomolecules 12:1045.

Machado Antonio, L., Henrique Martins, G., Zambon Barbosa Aragão, A., Galdi Quel, N., Zazeri, G., Houry, W. A., and Henrique Inacio Ramos, C. (2023). Unveiling the Role of Sorghum RPAP3 in the Function of R2TP Complex: Insights into Protein Assembly in Plants. Plants 12:2925.

Majerska, J., Schrumpfova, P. P., Dokladal, L., Schorova, S., Stejskal, K., Oboril, M., Honys, D., Kozakova, L., Polanska, P. S., and Sykorova, E. (2017). Tandem affinity purification of AtTERT reveals putative interaction partners of plant telomerase in vivo. Protoplasma 254:1547–1562.

Marchant, D. B., Chen, G., Cai, S., Chen, F., Schafran, P., Jenkins, J., Shu, S., Plott, C., Webber, J., Lovell, J. T., et al. (2021). Dynamic genome evolution in a model fern. Nat Plants 8:1038–1051.

Martino, F., Pal, M., Munoz-Hernandez, H., Rodriguez, C. F., Nunez-Ramirez, R., Gil-Carton, D., Degliesposti, G., Skehel, J. M., Roe, S. M., Prodromou, C., et al. (2018). RPAP3 provides a flexible scaffold for coupling HSP90 to the human R2TP co-chaperone complex. Nat Commun 9:3063.

Maurizy, C., Quinternet, M., Abel, Y., Verheggen, C., Santo, P. E., Bourguet, M., A, C. F. P., Bragantini, B., Chagot, M. E., Robert, M. C., et al. (2018). The RPAP3-C terminal domain identifies R2TP-like quaternary chaperones. Nat Commun 9:2093.

Muñoz-Hernández, H., Pal, M., Rodríguez, C. F., Fernández-Leiro, R., Prodromou, C., Pearl, L. H., and Llorca, O. (2019). Structural mechanism for regulation of the AAA-ATPases RUVBL1-RUVBL2 in the R2TP co-chaperone revealed by cryo-EM. Sci Adv 5:eaaw1616.

Nano, N., and Houry, W. A. (2013). Chaperone-like activity of the AAAþproteins RVb1 and RVb2 in the assembly of various complexes. Philosophical Transactions of the Royal Society B: Biological Sciences 368:1–12.

Pal, M., Morgan, M., Phelps, S. E., Roe, S. M., Parry-Morris, S., Downs, J. A., Polier, S., Pearl, L. H., and Prodromou, C. (2014). Structural basis for phosphorylation-dependent recruitment of Tel2 to Hsp90 by Pih1. Structure 22:805–818.

Punjani, A., Rubinstein, J. L., Fleet, D. J., and Brubaker, M. A. (2017). CryoSPARC: Algorithms for rapid unsupervised cryo-EM structure determination. Nat Methods 14:290–296.

Queitsch, C., Sangster, T. A., and Lindquist, S. (2002). Hsp90 as a capacitor of phenotypic variation. Nature 417:618–624.

Rivera-Calzada, A., Pal, M., Munoz-Hernandez, H., Luque-Ortega, J. R., Gil-Carton, D., Degliesposti, G., Skehel, J. M., Prodromou, C., Pearl, L. H., and Llorca, O. (2017). The Structure of the R2TP Complex Defines a Platform for Recruiting Diverse Client Proteins to the HSP90 Molecular Chaperone System. Structure 25:1145–1152 e4.

Roe, S. M., Prodromou, C., O’Brien, R., Ladbury, J. E., Piper, P. W., and Pearl, L. H. (1999). Structural basis for inhibition of the Hsp90 molecular chaperone by the antitumor antibiotics radicicol and geldanamycin. J Med Chem 42:260–266.

Samaras, P., Schmidt, T., Frejno, M., Gessulat, S., Reinecke, M., Jarzab, A., Zecha, J., Mergner, J., Giansanti, P., Ehrlich, H. C., et al. (2020). ProteomicsDB: A multi-omics and multi-organism resource for life science research. Nucleic Acids Res 48:D1153– D1163.

Sardiu, M. E., Cai, Y., Jin, J., Swanson, S. K., Conaway, R. C., Conaway, J. W., Florens, L., and Washburn, M. P. (2008). Probabilistic assembly of human protein interaction networks from label-free quantitative proteomics. Proc Natl Acad Sci U S A 105:1454–1459.

Schopf, F. H., Biebl, M. M., and Buchner, J. (2017). The HSP90 chaperone machinery. Nat Rev Mol Cell Biol 18:345–360.

Schorova, S., Fajkus, J., Zaveska Drabkova, L., Honys, D., and Schrumpfova, P. P. (2019). The plant Pontin and Reptin homologues, RuvBL1 and RuvBL2a, colocalize with TERT and TRB proteins in vivo, and participate in telomerase biogenesis. Plant J 98:195–212.

Serna, M., Gonz, A., Degliesposti, G., Zarzuela, E., Skehel, J. M., and Mu, J. (2021). CryoEM of RUVBL1 – RUVBL2 – ZNHIT2, a complex that interacts with pre-mRNA-processing-splicing factor 8. Nucleic Acids Res 50:1128–1146.

Sotta, N., Shantikumar, L., Sakamoto, T., Matsunaga, S., and Fujiwara, T. (2016). TPR5 is involved in directional cell division and is essential for the maintenance of meristem cell organization in Arabidopsis thaliana. J Exp Bot 67:2401–2411.

Van Leene, J., Eeckhout, D., Cannoot, B., De Winne, N., Persiau, G., Van De Slijke, E., Vercruysse, L., Dedecker, M., Verkest, A., Vandepoele, K., et al. (2015). An improved toolbox to unravel the plant cellular machinery by tandem affinity purification of Arabidopsis protein complexes. Nat Protoc 10:169–187.

Webb, B., and Sali, A. (2016). Comparative protein structure modeling using MODELLER. Curr Protoc Bioinformatics 2016:5.6.1–5.6.37.

Wing, Y., Cheung, S., Ramos, C. H. I., Kukura, P., and Houry, W. A. (2022). Assembly principles of the human R2TP chaperone complex reveal the presence of R2T and R2P complexes. Structure 30:1–16.

Yang, Z., Zeng, X., Zhao, Y., and Chen, R. (2023). AlphaFold2 and its applications in the fields of biology and medicine. Signal Transduct Target Ther 8:115.

Zhang, K. (2016). Gctf: Real-time CTF determination and correction. J Struct Biol 193:1– 12.

Zhao, R., Davey, M., Hsu, Y. C., Kaplanek, P., Tong, A., Parsons, A. B., Krogan, N., Cagney, G., Mai, D., Greenblatt, J., et al. (2005). Navigating the chaperone network: An integrative map of physical and genetic interactions mediated by the hsp90 chaperone. Cell 120:715–727.

Zheng, S. Q., Palovcak, E., Armache, J. P., Verba, K. A., Cheng, Y., and Agard, D. A. (2017). MotionCor2: Anisotropic correction of beam-induced motion for improved cryo-electron microscopy. Nat Methods 14:331–332.

Zhou, C. Y., Stoddard, C. I., Johnston, J. B., Trnka, M. J., Echeverria, I., Palovcak, E., Sali, A., Burlingame, A. L., Cheng, Y., and Narlikar, G. J. (2017). Regulation of Rvb1/Rvb2 by a Domain within the INO80 Chromatin Remodeling Complex Implicates the Yeast Rvbs as Protein Assembly Chaperones. Cell Rep 19:2033–2044.

Zivanov, J., Nakane, T., Forsberg, B. O., Kimanius, D., Hagen, W. J., Lindahl, E., and Scheres, S. H. (2018). New tools for automated high-resolution cryo-EM structure determination in RELION-3. Elife 7:e42166.

