## Supplementary Figures and Tables for "The structure of the R2T complex reveals a different architecture of the related HSP90 co-chaperones R2T and R2TP"

| | AGI Locus Code | Peptides | |
|---|---|---|---|
| | | R1 | R2 |
| **PFD1** | AT2G07340 | 90 | 107 |
| **PFD2** | AT3G22480 | 437 | 491 |
| **PFD3** | AT5G49510 | 498 | 530 |
| **PFD4** | AT1G08780 | 121 | 130 |
| **PFD5** | AT5G23290 | 205 | 228 |
| **PFD6** | AT1G29990 | 669 | 697 |

**Supplementary Figure S1.** Table indicating subunits of the PFD complex identified by TAP-MS using GS-PFD6 as bait. Number of unique peptides are intidated for each replicate (R).

A

| | AGI Code | Peptides | |
|---|---|---|---|
| | | R1 | R2 |
| **NRPB3** | AT2G15430 | 1 | 2 |
| **NRPB5A** | AT3G22320 | 18 | 15 |
| **NRPB5C** | AT5G57980 | 12 | 14 |
| **NRPB8B** | AT3G59600 | 4 | 3 |
| **NRPC1** | AT5G60040 | 19 | 13 |

B

| | AGI Locus Code | Peptides | | | Control | |
|---|---|---|---|---|---|---|
| | | R1 | R2 | R3 | R1 | R2 |
| **NRPA1** | AT3G57660 | 15 | 18 | 0 | 0 | 0 |
| **NRPB1** | AT4G35800 | 8 | 7 | 1 | 0 | 0 |
| **NRPB3** | AT2G15430 | 7 | 7 | 3 | 1 | 1 |
| **NRPB5C** | AT5G57980 | 4 | 9 | 4 | 0 | 0 |
| **NRPC1** | AT5G60040 | 48 | 23 | 3 | 0 | 0 |
| **NRPC2** | AT5G45140 | 2 | 1 | 0 | 0 | 0 |
| **NRPD2** | AT3G23780 | 2 | 2 | 1 | 1 | 0 |

**Supplementary Figure S2.** (A) Subunits of the nuclear RNA polymerases identified by TAP-MS using GS-PFD6. (B) Subunits of the nuclear RNA polymerases identified by AP-MS using GS-AtRPAP3 as bait. Number of unique peptides are intidated for each replicate (R).

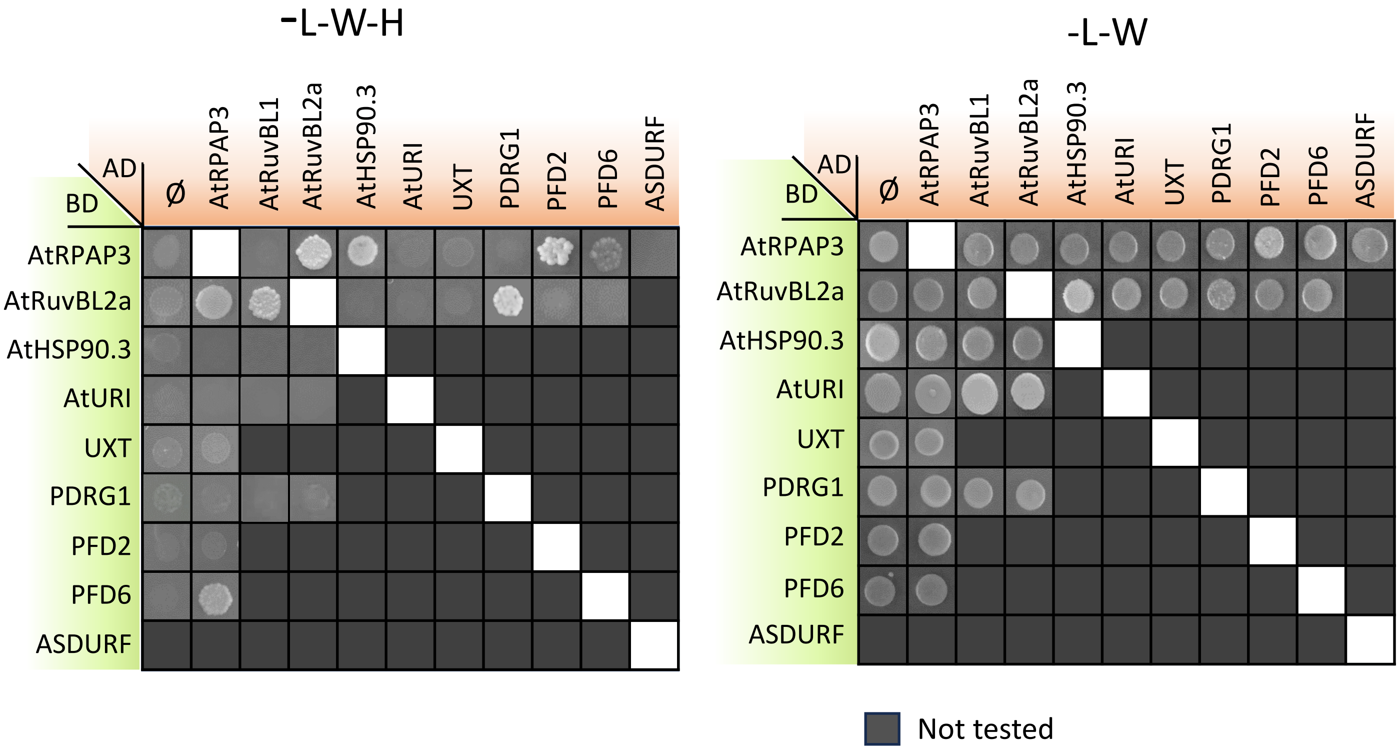


**Supplementary Figure S3.** Drop assay showing the interactions between subunits of R2T and PFDL complexes by Y2H. L, leucine; W, tryptophan; H, histidine.

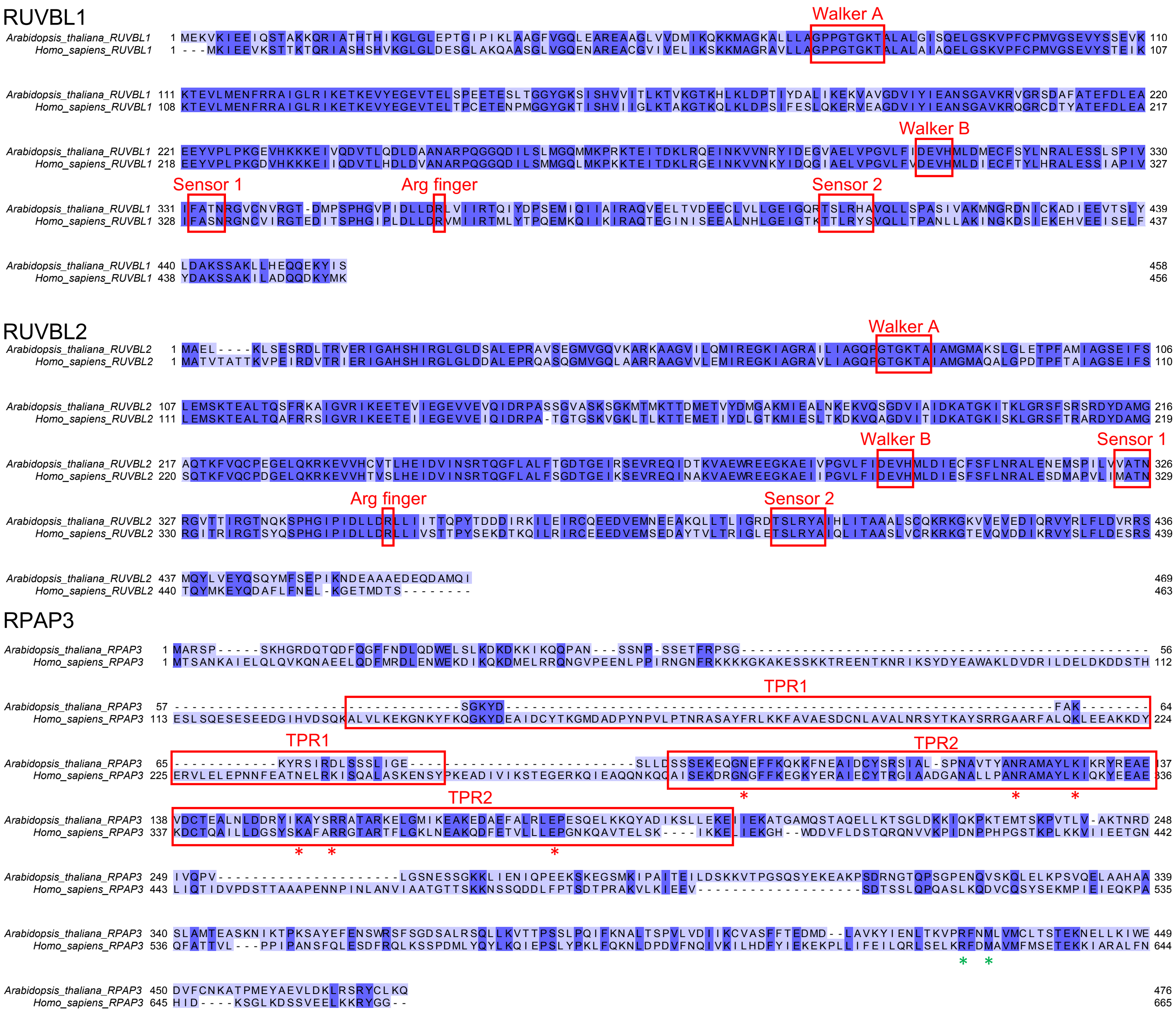


**Supplementary Figure S4.** Sequence alignment of RUuvL1, RuvBL2 and RPAP3 from *Homo sapiens* and *Arabidopsis*. The *Arabidopsis* RuvBL2 is AtRuvBL2a. Catalytic residues of RuvBL1 and RuvBL2 and the TPR domains of RPAP3 are indicated. Conserved amino acids (dark blue) were calculated with JALVIEW from the aligned sequences. Green asterisks mark the residues involved in the interaction between AtRPAP3 RBD and AtRuvBL2a.

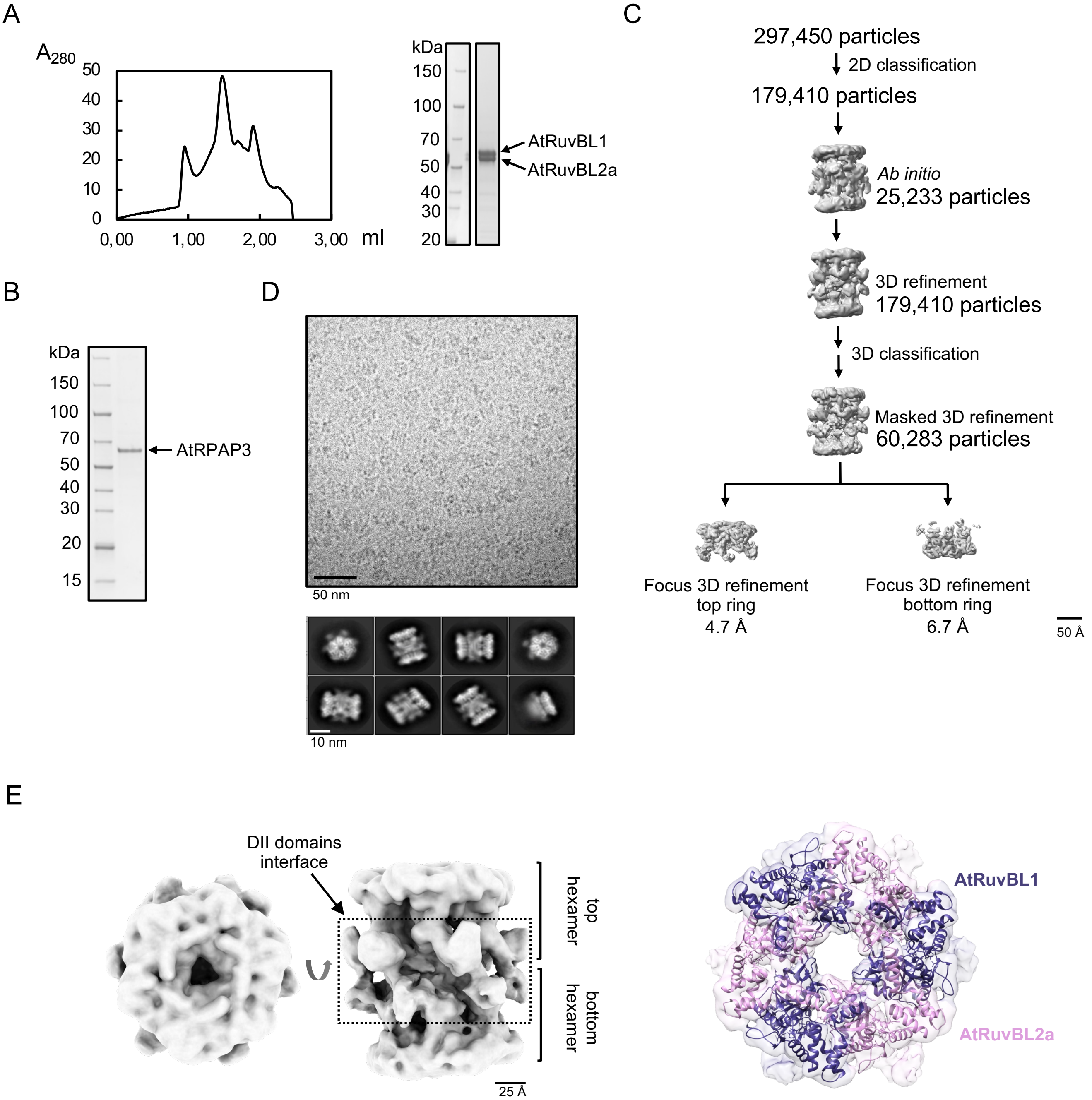


**Supplementary Figure S5.** Structural characterization of the *Arabidopsis* AtRuvBL1-AtRuvBL2a complex. (A) Size exclusion chromatography was used as a final step of purification for recombinant AtRuvBL1-AtRuvBL2a. Coomassie-stained SDS-PAGE shows the peak fraction from the chromatography used for structural experiments and ATPase activity. (B) Coomassie-stained SDS-PAGE of purified recombinant AtRPAP3 used in the ATPase activity of AtRuvBL1-AtRuvBL2a. (C) Image processing workflow of the cryo-EM images of the AtRuvBL1-AtRuvBL2a complex. Scale bar represents 50 Å. (D) Representative cryoEM field (top) and 2D averages (bottom) of AtRuvBL1-AtRuvBL2a. (E) CryoEM 3D reconstruction of the dodecameric AtRuvBL1-AtRuvBL2a complex highlighting the top and bottom heterohexameric modules and the interaction surface made up of DII domains. Right panel shows a top view of the complex with the atomic model of the human homologue rigid body fitted in the cryoEM density (PDB 2XSZ). Scale bars are indicated.

A

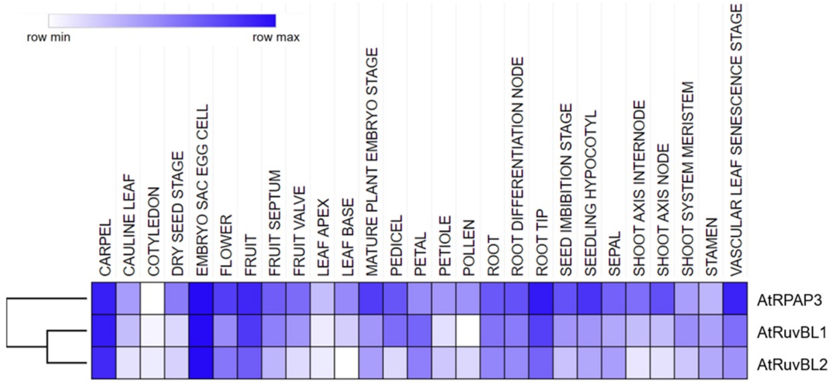


B

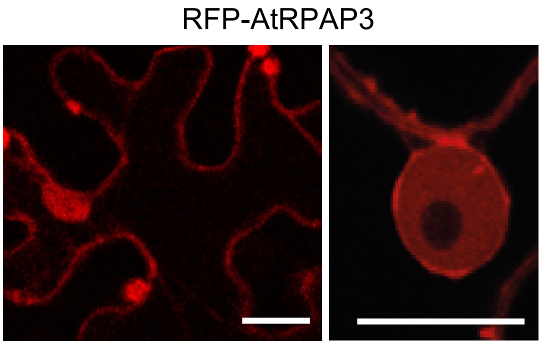


**Supplemental Figure S6.** (A) Normalized protein accumulation in various *Arabidopsis* organs using ProteomicsDB. AtRuvBL2 refers to AtRuvBL2a. (B) Nuclear and cytoplasmic localization of AtRPAP3, fused to RFP tag, in *N. Benthamina* leaves (Scale bar: 10µm).

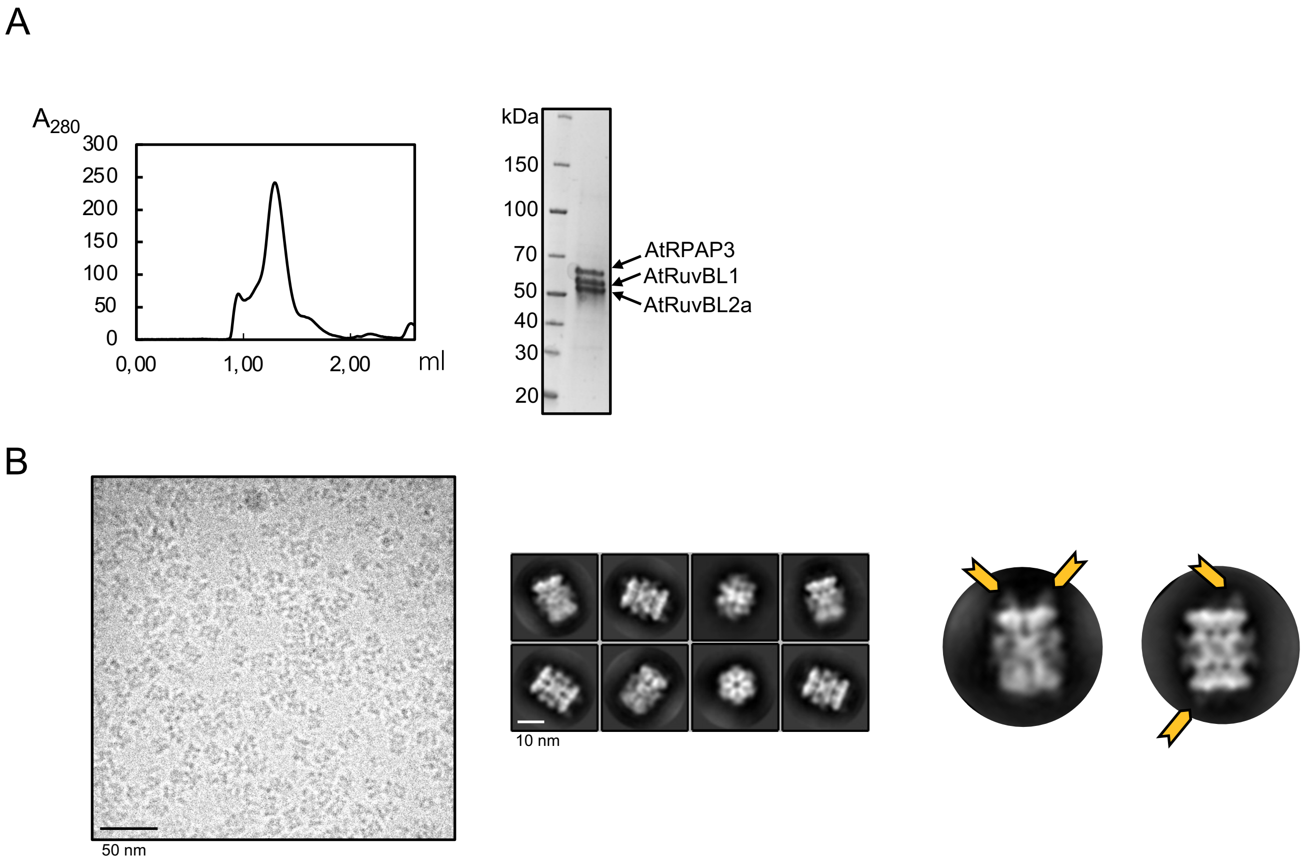


**Supplementary Figure S7.** Purification and preliminary structural characterization of the *Arabidopsis* R2T complex. (A) AtRuvBL1, AtRuvBL2a and AtRPAP3 where co-expressed to favor complex formation *in vivo*. Chromatogram shows the size exclusion purification of ther R2T complex. Coomassie-stained SDS-PAGE shows the peak fraction from the chromatography used cryo-EM experiments. (B) CryoEM field (left) and 2D averages (middle) of R2T. Scale bars are indicated. The complex displayed a dodecameric architecture similar to AtRuvBL1-AtRuvBL2a with extra densities attached to the AAA+ face of the hexameric rings (labeled with yellow arrows in the right inserts).

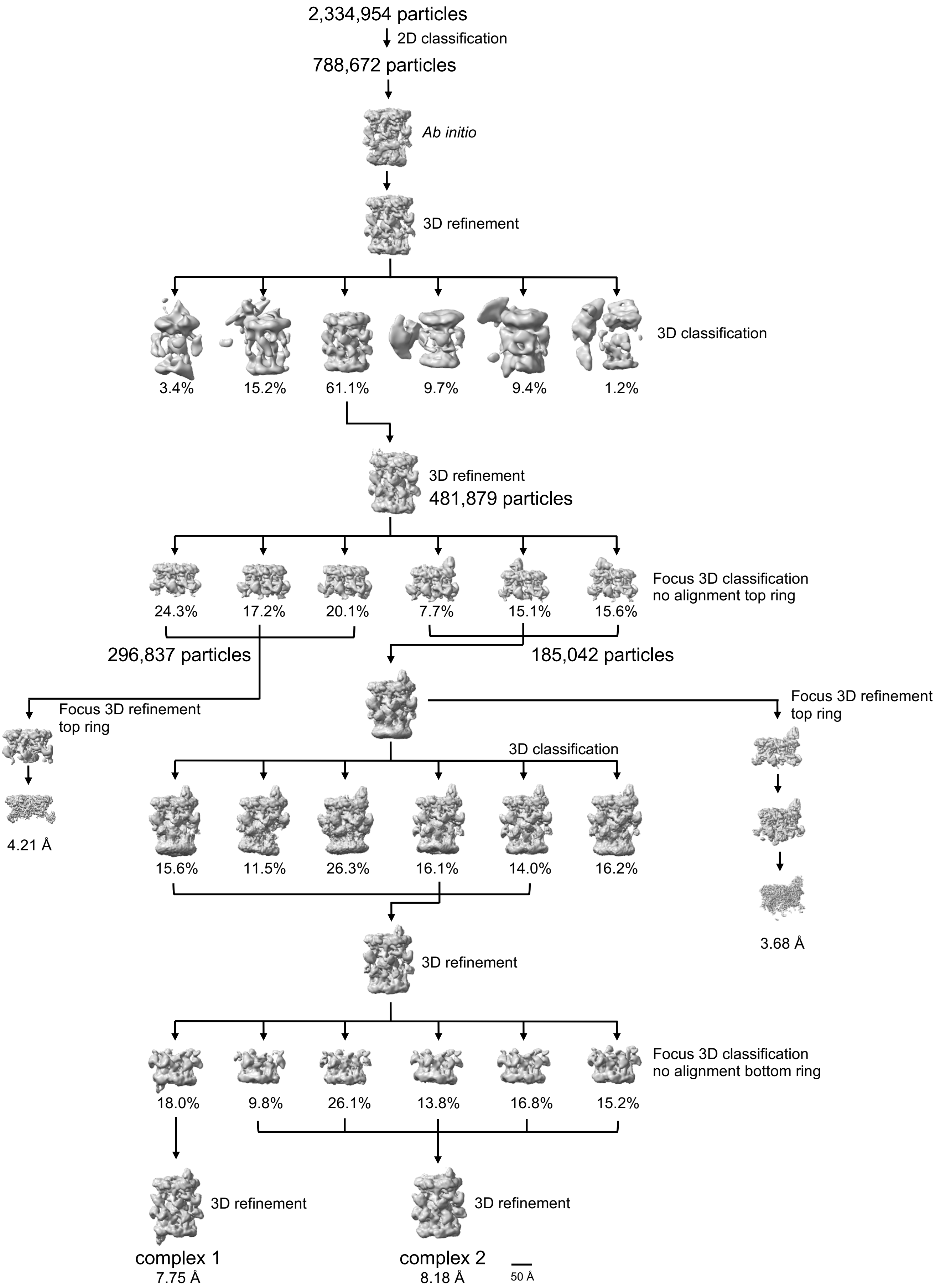


**Supplementary Figure S8**. Image processing workflow of the cryoEM images of the R2T complex. Scale bar represents 50 Å.

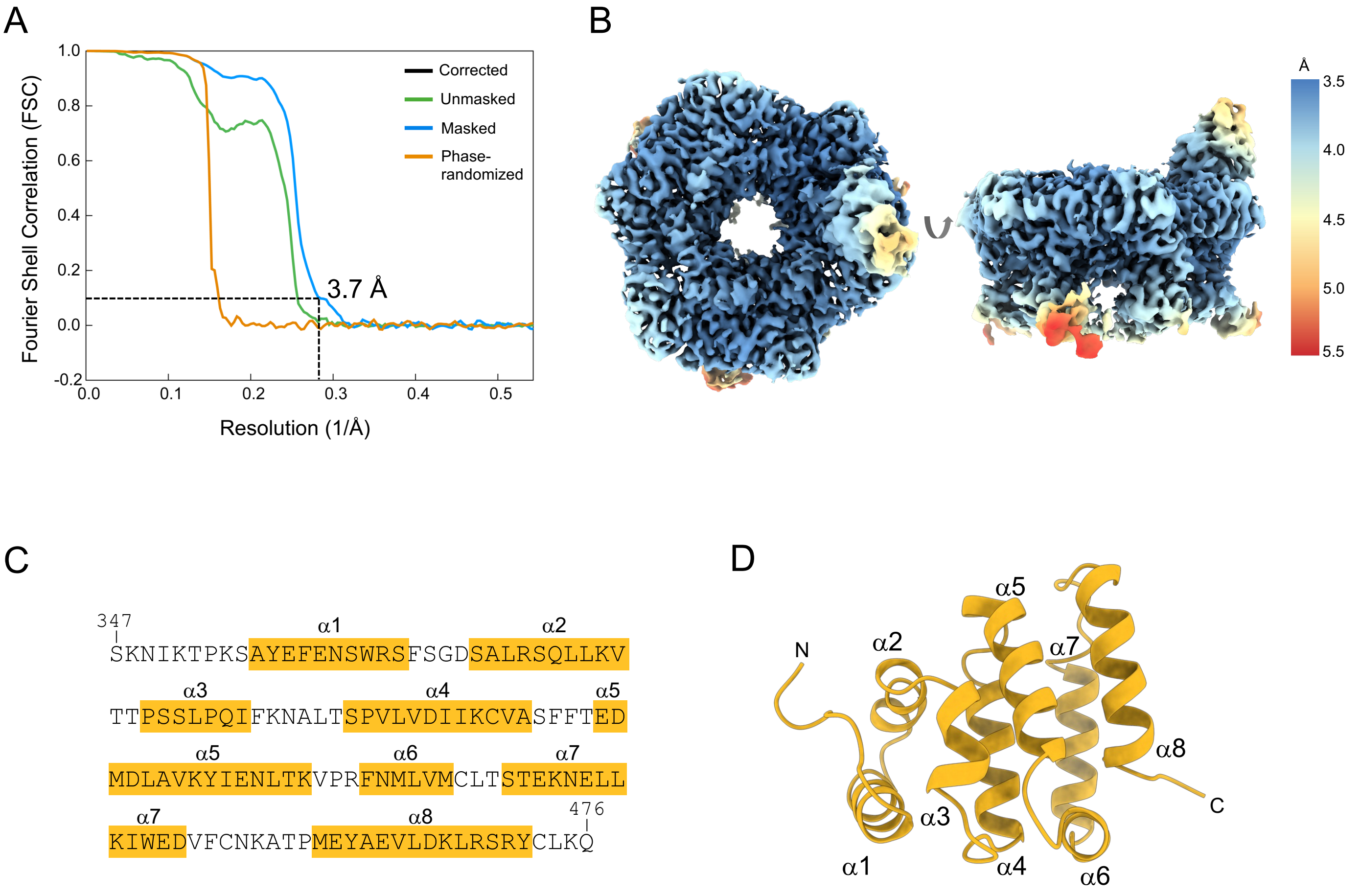


**Supplementary Figure S9.** Resolution estimation of the cryoEM for the R2T top ring. (A) Fourier Shell Correlation (FSC) curves for the R2T core complex obtained by focus refinement of the top ring in the dodecameric structure. (B) Local resolution estimates for the R2T top ring complex as provided by RELION. Bottom and side views of the map are shown using the color scale shown on the right. (C) Sequence of AtRPAP3 RBD domain highlighting the α-helices in orange. (D) Ribbon representation of the AtRPAP3 RBD domain from the R2T complex. α-helices 1 to 8 are indicated.

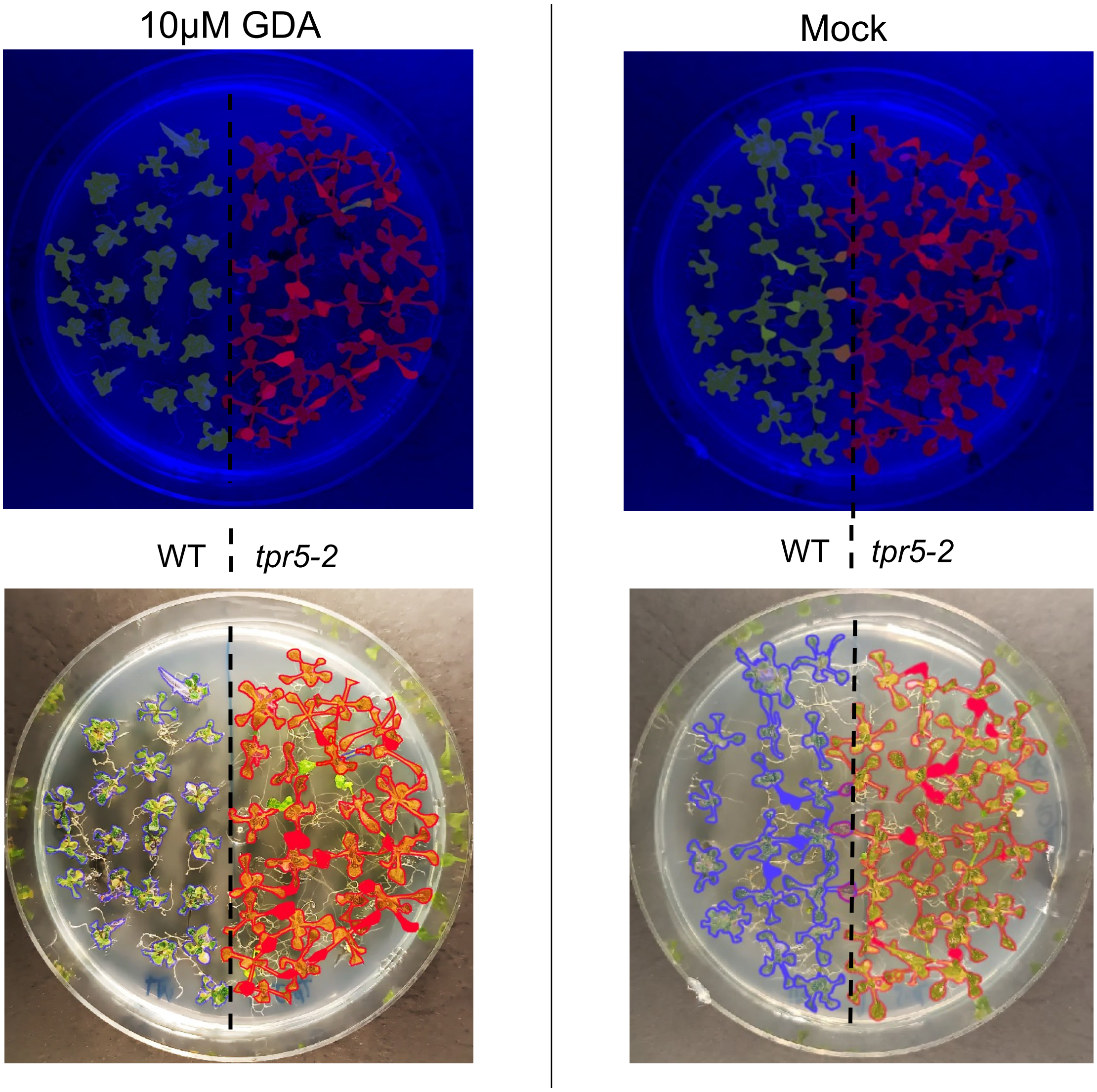


**Supplementary Figure S10.** Recognition of seedlings with BioDock. Two classes corresponding to WT and the *tpr5-2* seedlings were defined for the images of each condition. First, an automatic detection of objects was performed for both classes (top), followed by a manual curation (bottom).

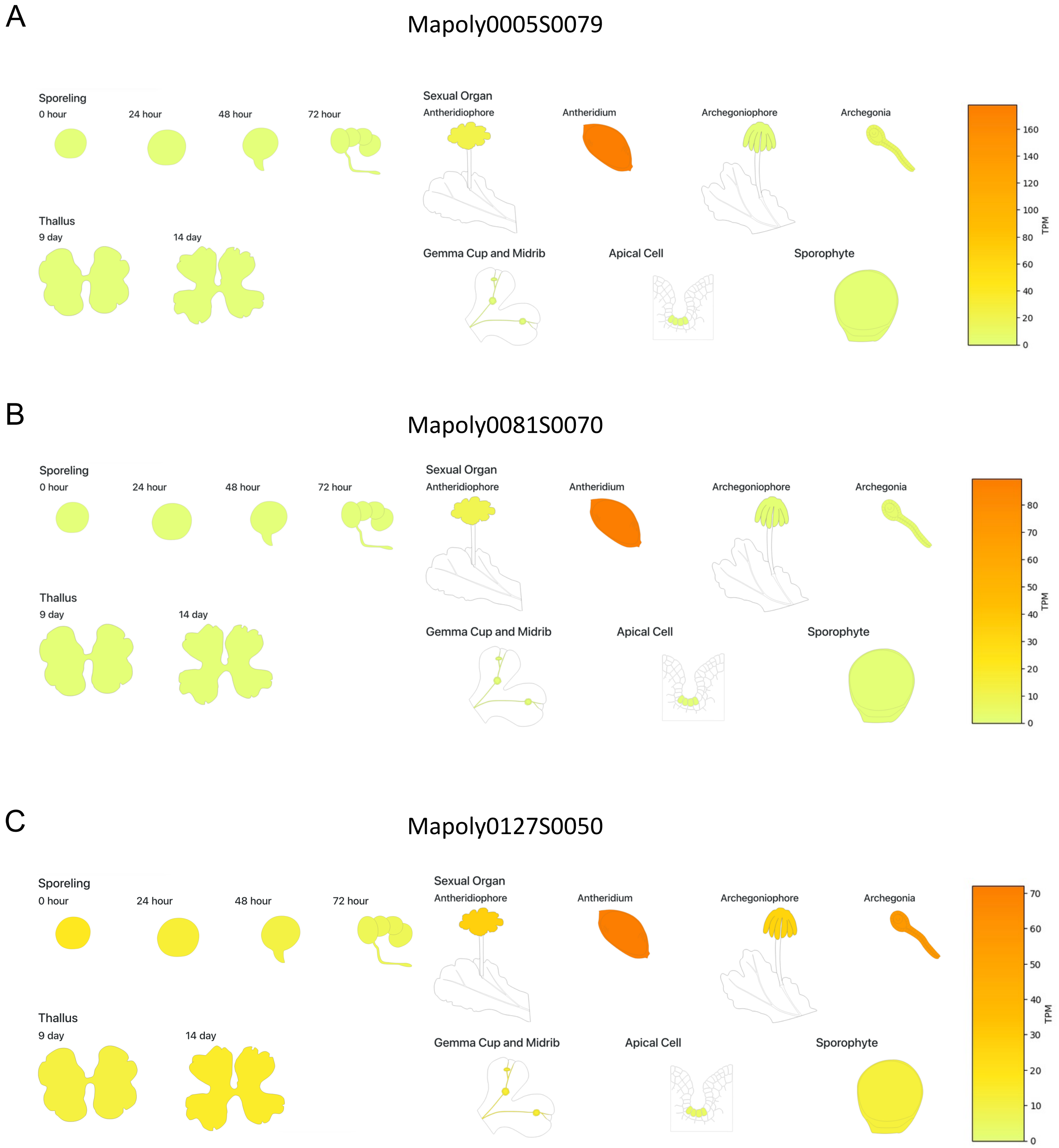


**Supplementary Figure S11.** Tissue-specific expression in *Marchantia polymorpha*. (A, B) Chromatic expression images of the two *Marchantia* genes encoding proteins with a PIH domain. (C) Chromatic expression images of the only gene of *Marchantia* encoding a protein with a TPR domain and an RBD found in RPAP3. Expression levels are shown in TPM in the color-coded bar. The data was obtained using the Expression Database for *Marchantia polymorpha* (https://marchantia.info/mbex/).

**Supplementary Table S1.** CryoEM data collection, refinement, and validation statistics.

| Structure | Arabidopsis thaliana R2T complex (EMD-19894) (PDB ID 9EQ2) |
|---|---|
| **Data collection and processing** | |
| Microscope | Titan Krios |
| Detector | Falcon 4i |
| Nominal magnification | 130,000 |
| Voltage (kV) | 300 |
| Electron exposure ($e^-/Å^2$) | 50.12 (50 fractions) |
| Defocus range (µm) | -1.5 to -3.0 |
| Pixel size (Å) | 0.921 |
| Symmetry imposed | C1 |
| Initial particle images (no.) | 2,334,954 |
| Final particle images (no.) | 185,042 |
| Map resolution (Å) | 3.68 |
| FSC threshold | 0.143 |
| Map resolution range (Å) | 3.5 – 5-5 |
| **Refinement** | |
| Initial model used | AlphaFold |
| Map sharpening B factor ($Å^2$) | -128.2 |
| **Model composition** | |
| Non-hydrogen atoms | 14628 |
| Protein residues | 1876 |
| Nucleotide residues | 0 |
| Ligands | ADP:3 |
| **R.m.s. deviations** | |
| Bond lengths (Å) | 0.007 |
| Bond angles (°) | 1.030 |
| **Validation** | |
| MolProbity score | 1.22 |
| Clashscore | 2.79 |
| Rotamers outliers (%) | 0.25 |
| **Ramachandran (%)** | |
| Favored | 97.17 |
| Allowed | 2.83 |
| Disallowed | 0.00 |
| **Model vs Data** | |
| CC volume | 0.72 |

**Supplementary Table S2.** Oligonucleotides used in this work.

| | |
|---|---|
| *Cloning of CDSs (Gateway)* | |
| AtRPAP3_attB1_Fw | GGGGACAAGTTTGTACAAAAAAGCAGGCTTCATGGCTAGGTCACCGAGCAA |
| AtRPAP3_attB2_Rv | GGGGACCACTTTGTACAAGAAAGCTGGGTTTACTGTTTAAGGCAGTATC |
| AtRPAP3_attB2_noSTOP_Rv | GGGGACCACTTTGTACAAGAAAGCTGGGTCCTGTTTAAGGCAGTATCTTGAC |
| AtRuvBL1_attB1_Fw | GGGGACAAGTTTGTACAAAAAAGCAGGCTTCATGGAGAAAGTAAAGATTGA |
| AtRuvBL1_attB2_Rv | GGGGACCACTTTGTACAAGAAAGCTGGGTCTGAGATGTATTTTTCTTGTT |
| AtRuvBL2a_attB1_Fw | GGGGACAAGTTTGTACAAAAAAGCAGGCTTCATGGCGGAACTAAAGCTATC |
| AtRuvBL2a_attB1_Fw | GGGGACCACTTTGTACAAGAAAGCTGGGTCGATCTGCATAGCATCTTGTT |
| AtRPAP3mut2_int_Fw | ACAAAGGTTCCTGCATTCAACGCACTCGTCATGTGCC |
| AtRPAP3mut2_int_Rv | GGCACATGACGAGTGCGTTGAATGCAGGAACCTTTGT |
| *Cloning in bacterial expression plasmids* | |
| pET15b_AtRuvBL1_BamHI_Fw | ATATGCTCGAGGATCCAATGGAGAAAGTAAAGATTGAAGAAATTCAGTCCA |
| pET15b_AtRuvBL1_BamHI_Rv | GTTAGCAGCCGGATCCTCATGAGATGTATTTTTCTTGTTGCTCATGC |
| pCDFDuet_AtRuvBL2a_KpnI_Fw | CGGGGTACCATGGCGGAACTAAAGCTAT |
| pCDFDuet_AtRuvBL2a_KpnI_Rv | CGGGGTACCTCAGATCTGCATAGCATCTTG |
| pRSFDuet_1_Cons_TEV_AtRPAP3_FW | GAGAGAATCTTTATTTTCAGGGCATGGCTAGGTCACCGAGCAAACACG |
| pRSFDuet_2_AtRPAP3_cons_3C_RV | TGGAACAGCACCTCCAGCTGTTTAAGGCAGTATCTTGACCTTAGTTTGTC |
| pRSFDuet_Vector_Cons_Fw | CTGGAGGTGCTGTTCCAGGGACCT |
| pRSFDuet_Vector_Cons_Rv | GCCCTGAAAATAAAGATTCTCTCCGCC |
| *Preparing the AtHSP90.3:6xHis-TEV-3xFLAG-AtHSP90.3 construct* | |
| -1141_TSS_AtHSP90.3_Fw | GGGACAAGTTTGTACAAAAAAGCAGGCTTCGGGGGAAAAGTGTGTAATATATAGGGGATG |
| AtHSP90.3_genomic_Rv | GGGGACCACTTTGTACAAGAAAGCTGGGTCTTAGTCAACTTCCTCCATCTTGCT |
| NcoI_6xHis-3xFlag_Fw | CATGCCATGGGTCATCACCATCACC |
| NcoI_6xHis-3xFlag_Rv | CATGCCATGGTTCCTTTGTCGTCA |
| AtHSP90-3flag_NcoI_Fw | CGATCAACGACCATGGGTCATCACCATCACCATCACGATTATGAT |
| AtHSP90-3flag_NcoI_Rv | TCCGTCGCGACCATGGTTCCTTTGTCGTCATCATCTTTATAGTCTTTATC |
